# MEA-LINK identifies the CCL4-CCR5 axis in neuronal hyperactivity control by human microglia

**DOI:** 10.64898/2026.02.04.703799

**Authors:** Annika Mordelt, Nicky Scheefhals, Imke M. E. Schuurmans, Koen Slottje, Mariela Cano Rivera, Marina P. Hommersom, Ao Huang, Leon Bichmann, Elly I. Lewerissa, Eline J. H. van Hugte, Dirk Schubert, John S. Tsang, Nael Nadif Kasri, Lot D. de Witte

**Affiliations:** Department of Human Genetics, Radboud university medical center, Nijmegen, the Netherlands.; Department of Medical Neuroscience, Donders Institute for Brain, Cognition and Behaviour, Radboud University, Nijmegen, the Netherlands; Donders Centre for Neuroscience, Donders Institute for Brain, Cognition and Behaviour, Radboud University, Nijmegen, the Netherlands; Center for Systems and Engineering Immunology and Department of Immunobiology, Yale University School of Medicine, New Haven, CT, USA; Chan Zuckerberg Biohub New York, New Haven, CT, USA; Department of Psychiatry, Radboud university medical center, Nijmegen, the Netherlands; Department of Psychiatry, Icahn School of Medicine at Mount Sinai, New York, NY 10029, USA

## Abstract

Microglia, the resident immune cells of the brain, act along a spectrum to maintain CNS homeostasis, respond to perturbations, and control neuronal activity. Disentangling the molecular mechanisms of human microglia-neuron crosstalk remains challenging due to the context-dependent, dynamic nature of their interaction. We introduce MEA-LINK, a systems-approach leveraging natural variation to screen for immune modulators of neuronal activity. This multi-modal platform integrates human induced pluripotent stem cell (hiPSC) technology with micro-electrode array (MEA) recordings and proteomic analyses of secreted immune factors, allowing for longitudinal samples and correlations across modalities. We applied MEA-LINK to explore microglia-neuron interactions during development and hyperactivity challenges. We show that human microglia accelerate neuronal network development and rescue hyperactive network phenotypes. Linking the secretome adaptations to neuronal network activity variations, we identified CCL4 as a top candidate in microglia-mediated hyperactivity control. Then, we functionally validated the context-dependent role of microglial CCL4 to neuronal CCR5 signaling in human neuronal networks. Our findings support a neuron-specific function of chemokines and their receptors in the brain and provide a new perspective for immune signaling in neuronal hyperactivity control. The MEA-LINK platform thus offers a foundation for comprehensive, systematic studies of human microglia-neuron interactions.

**Highlights:**

- MEA-LINK integrates micro-electrode array recordings with proteomics of longitudinal samples to identify immune modulators of neuronal activity.
- Human microglia rescue neuronal hyperactivity induced by pharmacological and genetic challenges.
- Microglial CCL4 dampens neuronal activity via CCR5 signaling in a context-dependent manner.

**Graphical abstract:** 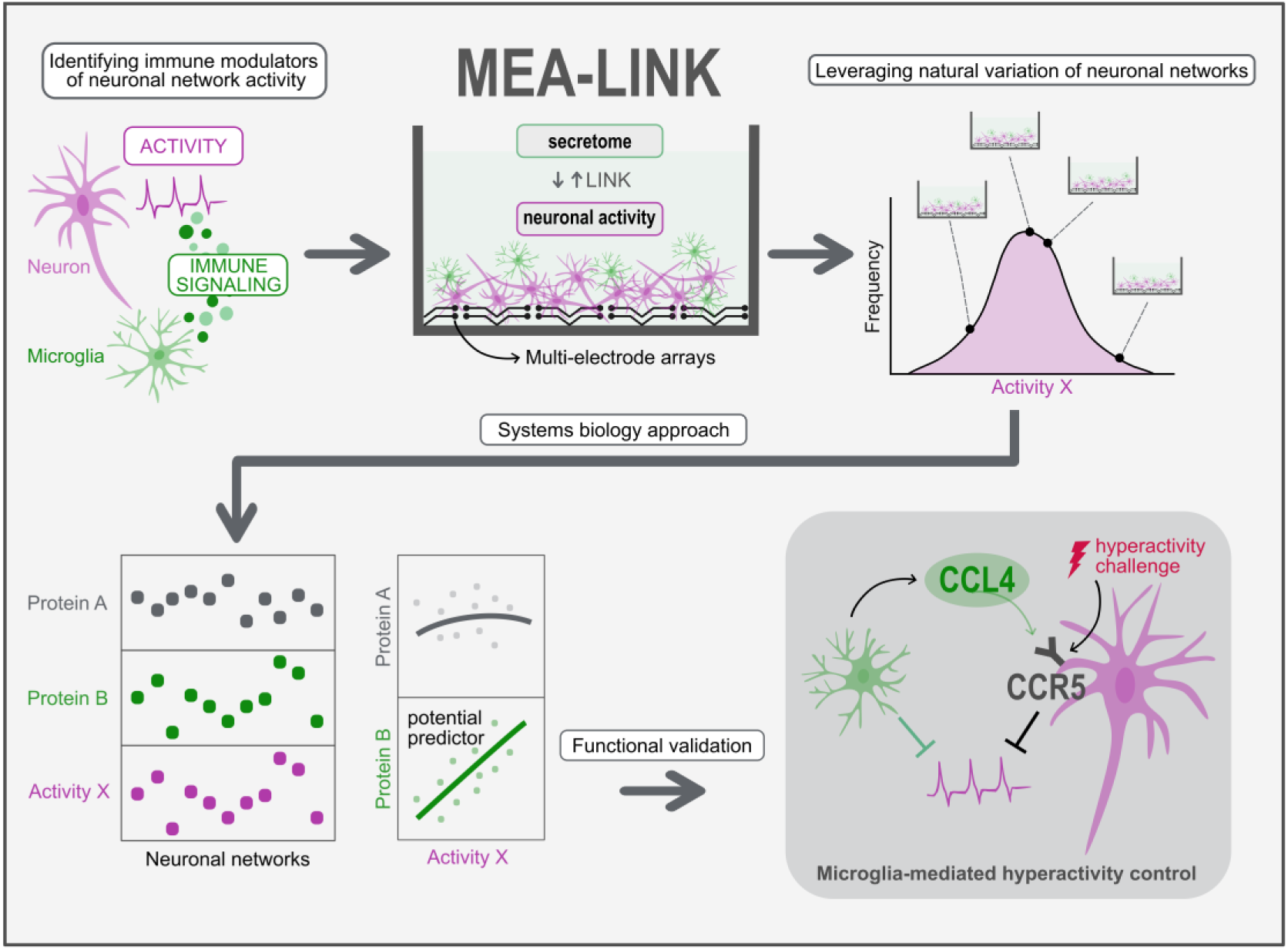

## Introduction

The immune system and the central nervous system (CNS) converge in many evolutionary advantageous principles. Both systems are constantly sensing and responding to environmental factors, mounting a behavioral or inflammatory response to ensure healthy brain function. Shared biological principles are experience-driven fine tuning^1, 2^, memory function^3, 4^, as well as homeostatic control^5, 6^. Maintaining homeostatic levels of neuronal activity and inflammation requires compensatory adjustments when deviations from baseline occur^7^. We hypothesize that homeostatic mechanisms of the CNS and immune system share common molecular machinery.

Microglia, the brain’s resident myeloid cells, represent a key link between the immune system and neuronal network formation and function. Unlike other macrophages, microglia have highly dynamic processes that constantly survey and contact nearby dendrites, axons and synapses^8–10^. They respond rapidly to environmental changes^11, 12^, including not only inflammatory cues like infection or injury but also shifts in neuronal activity^13^. This bidirectional microglia-neuron communication is thought to play an important role in brain function, and dysfunctional microglia have been associated with a variety of CNS disorders^14–16^. How human microglia and their immune signaling in turn influences neuronal network activity remains largely unclear.

Disentangling the molecular mechanisms of this microglia-neuron crosstalk has proven to be challenging, as it involves complex cell-cell interactions that must be robustly represented in the model system used. Further, many experimental models present extreme cases outside the physiological range (e.g. skewing microglial phenotypes with for instance Lipopolysaccharide (LPS) stimulation or complete gene knockouts). It is now widely acknowledged that immune cells, including microglia, act along a dynamic spectrum, and that their functions cannot be categorized in a dichotomous manner^17^. Building on approaches from the systems immunology field^18, 19^, our solution for studying microglia-neuron crosstalk beyond extreme cases is to utilize natural variation. Biological quantities like genes and proteins are inherently stochastic^20^, and these variations can be leveraged to establish correlations among parameters from different modalities^21^. We introduce MEA-LINK: a systems approach for studying functional microglia-neuron interactions in a human context. This multi-modal platform integrates longitudinal micro-electrode array (MEA) recordings with paired proteomic analyses of human induced pluripotent stem cell (hiPSC)-derived microglia-neuron co-cultures.

Using MEA-LINK, we screened for microglia-dependent immune mediators that regulate neuronal activity during development and in hyperactivity challenges. We chose these applications as there is a strong interest in the role of microglia in neurodevelopment^9, 22, 23^ and in the context of epilepsy^24^, where microglia were initially thought to be harmful but are now considered potentially neuroprotective^25, 26^. For example, Badimon et al. showed that rodent microglia can suppress excessive neuronal activity through a feedback mechanism resembling that of inhibitory neurons^27^. Consistently, removing microglia, pharmacologically or genetically, aggravated seizures in several rodent models^25^. However, microglial functions and glia-neuron signaling differ across species^12, 28^, and it remains unclear whether similar microglia-mediated protection occurs in the human brain. Post-mortem tissue from epilepsy patients shows increased inflammation^29, 30^, but how this relates to a protective microglial role is still unresolved. Here, we applied MEA-LINK to seizure-like conditions to identify immune mediators of neuronal activity regulation by human microglia.

## Results

### Human microglia enhance neuronal network synchrony and reduce duration of network bursts early in development

Emerging hiPSC technologies have advanced models of human brain cells substantially, including protocols to generate both neurons and microglia^31–35^. To study the complex and continuous interplay between these two cell types, a long-term co-culture of neurons and microglia is required (Fig. 1a). We previously developed an optimized co-culture model where both hiPSC-derived microglia and glutamatergic neuronal networks differentiate alongside each other allowing for cell-cell contact during maturation^36^. The engineered co-culture environment induces the typical ramified microglial morphology and expression of microglial signature markers^36^. To analyze neuronal network activity, we recorded spontaneous activity of neurons using micro-electrode arrays (MEAs). With increasing number of days *in vitro* (DIV) neuronal networks followed a robust developmental trajectory and became increasingly organized showing synchronized bursts (Suppl. Fig. 1a). This was reflected by an increase in mean firing rate (MFR), synchrony and network burst percentage (NBP), and a decrease in network burst duration (NBD) (Fig. 1b), as shown previously^37^. The expression of gene sets linked to synaptic function in human glutamatergic neurons^38^ showed strong correlations with the neuronal network activity parameters from the same network (Suppl. Fig. 1e), showing that increased network organization mirrors maturation of synaptic transmission. The directionality of the correlation was in line with the change during development, with synchrony showing positive correlations with the synaptic genes, whereas NBD was negatively correlated (Suppl. Fig. 1f).

**Figure 1.**
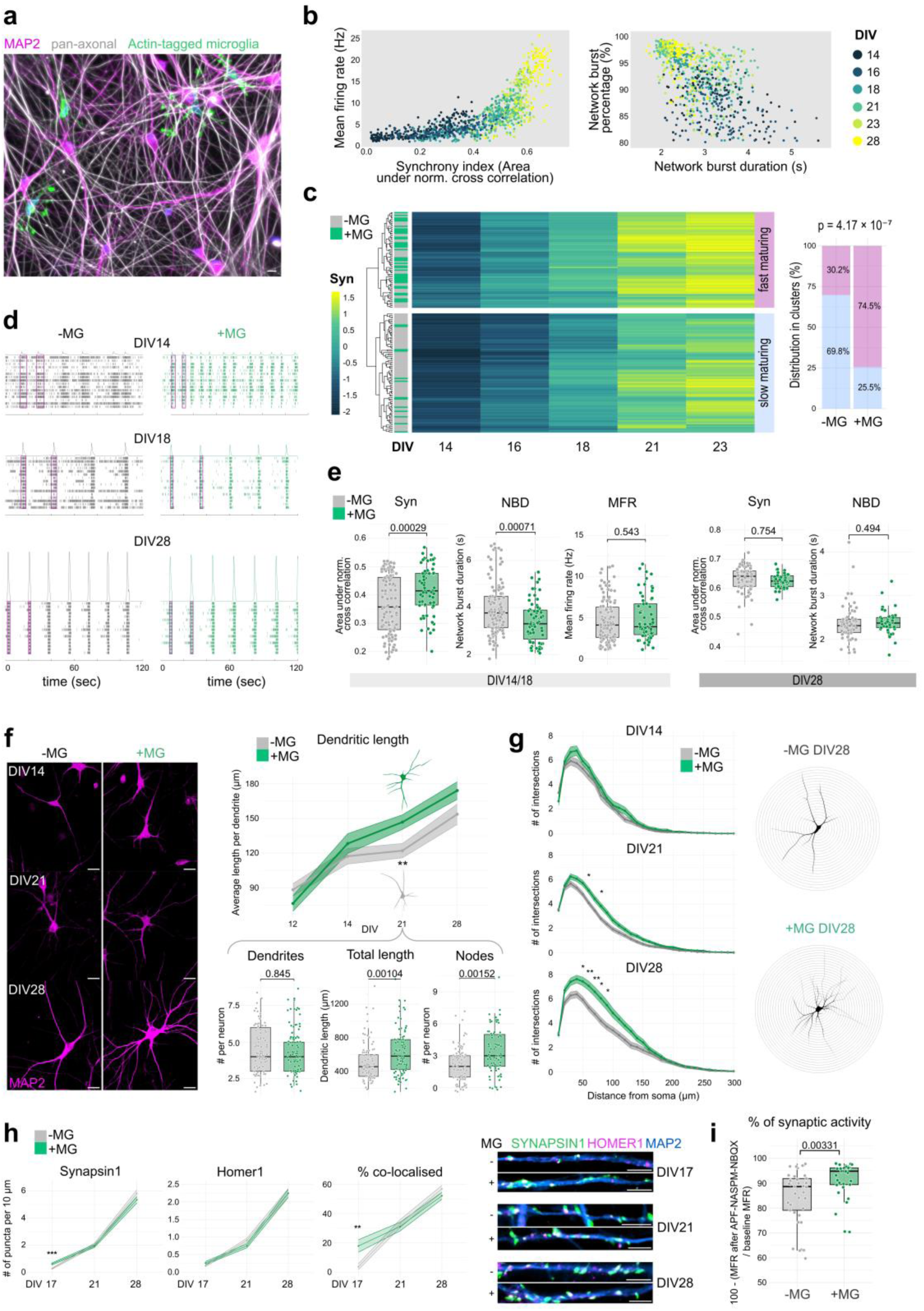
Human microglia accelerate the maturation of neuronal networks. **(a)** Representative fluorescence microscope image of hiPSC-derived microglia-neuron co-culture, scale bar = 10 μm. **(b)** Dot plots of neuronal networks color coded for DIV showing synchrony (Area under normalized cross-correlation) vs. mean firing rate (Hz) (top) and network burst duration (sec) vs. network burst percentage, N = 4, n = 1,071 neuronal network recordings. **(c)** Heatmap using unsupervised Euclidean clustering of neuronal networks based on synchrony across developmental timeline, N = 4, n = 157 neuronal networks each recorded at 5 timepoints (left), and distribution of clusters per condition with Chi-square test (right), MG = microglia. **(d)** Representative raster plots of neuronal network activity per condition at DIV14, DIV18, and DIV28. **(e)** Quantification of synchrony (DIV18 and DIV28), network burst duration (DIV14 and DIV28), and mean firing rate (DIV18) per condition, N = 4, n = 112 neuronal networks for -MG, n = 63 neuronal networks for +MG, unpaired t-test. **(f)** Representative fluorescence microscope images of MAP2-labeled neurons at DIV14, DIV21, and DIV28 per condition, scale bar = 20 μm (left) and quantification of neuronal morphology showing average length per dendrite (μm) across development, N = 3, n = 222 neurons for -MG, n = 211 for +MG, unpaired-test with post-hoc Bonferroni correction, **p <0.01 (top right) and the number of dendrites, total dendritic length, and nodes at DIV21, n = 93 neurons for -MG, n = 93 neurons for +MG, unpaired t-test (bottom right). **(g)** Sholl Analysis showing number of intersections in regard to the distance to the soma at DIV14, DIV21, and DIV28, *p <0.05, **p <0.01, unpaired t-test with post hoc Bonferroni correction (left) and representative Scholl analysis image based on somatodendritic reconstruction at DIV28 per condition (right). **(h)** Quantification of Synapsin1-, Homer1b/c-, and co-localized puncta per 10 μm dendrite at DIV17, DIV21, and DIV28, N = 2, n = 388 dendrites for -MG, n = 428 dendrites for +MG, **p <0.01, ****p <0.0001, unpaired t-test with post hoc Bonferroni correction (left) and representative fluorescence microscope images, scale bars = 5 μm (right). **(i)** Quantification of the percentage of neuronal activity lost after pharmacological blocking with AP5, NASPM, and NBQX, N = 3, n = 37 neuronal networks for -MG, n = 32 neuronal networks for +MG, unpaired t-test.

To investigate the role of microglia on neuronal network development, we compared neuronal network activity in networks without (-MG) or with microglia (+MG) in four batches across their developmental trajectory. We approached the resulting dataset consisting of >1,000 individual network recordings in an unsupervised manner. Euclidean clustering analysis of the synchrony index per well across the developmental trajectory revealed two distinct clusters (Fig. 1c). The upper cluster showed higher synchrony at earlier timepoints, indicating faster network maturation. When comparing the distributions, +MG networks were significantly enriched in the fast-maturing cluster. Indeed, group-wise comparisons early in development showed a significant increase in synchrony in +MG networks (Fig. 1d-e). We consistently observed this phenotype in the early DIVs in all four batches (Suppl. Fig. 2e). Furthermore, NBD was decreased in +MG networks early in development, whereas the MFR remained unchanged, suggesting a reorganization of network activity rather than merely increase in activity (Fig. 1e). By DIV28, differences in synchrony and NBD were no longer present, consistent with faster maturation rather than permanent changes in network activity per se. Interestingly, this microglia-mediated acceleration of network activity maturation was enhanced when doubling the amount of microglia added to the cultures, suggesting a dose-dependent effect (Suppl. Fig. 2f). To rule out potential confounding factors, we recorded neuronal activity before the addition of the microglia at DIV9 and observed no pre-existing differences in network organization (Suppl. Fig. 2b). Furthermore, culture viability at DIV28 was comparable between conditions (Suppl. Fig. 2c), and the top upregulated genes in +MG versus -MG networks were exclusively microglia-associated markers, verifying successful microglia integration and survival in the MEA wells (Suppl. Fig. 2d).

### Human microglia promote dendritic growth and synaptic network maturation

Given the accelerated development of neuronal network activity in the presence of microglia, we next investigated dendritic growth and arborization across development (Fig. 1f). Somatodendritic reconstructions revealed increased dendritic length in +MG networks starting at DIV21. The total number of dendrites was not different at DIV21, but the number of nodes per neuron and the total dendritic length were significantly increased in +MG networks, indicating that the existing dendrites extended further and arborized more. Sholl analysis confirmed more complex dendritic branching in +MG networks at DIV21 and DIV28 (Fig. 1g). Next, we explored whether this enhanced dendritic growth affected synapse development. Quantification of Homer1b/c- and Synapsin1-puncta along dendrites showed the expected increase in synaptic density across the developmental timeline (Fig. 1h). The +MG networks showed a significantly higher Synapsin1 density at DIV17 and then converged with the -MG networks at later time points, paralleling the development of neuronal network activity (Fig. 1d). Although the Homer1b/c density did not differ between conditions, the percentage of Homer1b/c co-localizing with Synapsin1 was significantly increased in +MG networks at DIV17 (Fig. 1h). Overall, the increased average dendrite length (Fig. 1f) and the comparable synapse density at later timepoints indicate a higher total synapse number in +MG networks compared to -MG networks. To determine whether this contributes functionally to neuronal network activity and maturation, we next assessed the dependence of neuronal network activity on synaptic transmission. MEA-recorded network activity represents both intrinsically generated action potentials and synaptically driven firing. By pharmacologically inhibiting AMPARs and NMDARs, we isolated the intrinsic, non-synaptically driven component of activity^39^. +MG networks showed a stronger reduction in firing upon glutamatergic receptor blockade, indicating that their activity was more dependent on synaptic transmission and therefore functionally more mature (Fig. 1i). In conclusion, human microglia enhanced dendritic growth and synaptic maturation of human neurons during development.

### Leveraging stochasticity to gain biological insights with MEA-LINK

Next, we explored potential mechanisms that could underlie the neuronal network-maturing function of human microglia. We hypothesized that this might be mediated via neuro-immune interactions, with secreted factors as one potential route of action. Secreted proteins, like cytokines, chemokines and growth factors, are well known for orchestrating communication between immune cells in the periphery to regulate inflammatory responses^40^. Recently, there has been increasing evidence that these molecules also modulate neuron-glia interactions^41–43^. Yet, the specific factors responsible for particular processes and the consequences of these interactions for neuronal development and function are still largely unknown.

Comparing MEA parameter activity distributions (Fig. 2a) illustrated the accelerated development of neuronal networks containing microglia and also highlighted the variability across individual replicates, a common feature of MEA data from cultured neurons^44^. Especially early in development, individual networks followed divergent trajectories, with some networks being already more mature than others (Fig. 2a). Later in development at DIV28, this variability decreased and networks converged toward a similar functional endpoint (Fig. 2a). Such variability is a prevalent feature of biological systems and is often underappreciated in experiments, as the focus is on the average behavior. These natural variations can in fact be leveraged to investigate the correlative relationship among system parameters. The correlations among parameters, particular those between different types of parameters such as neuronal activity and secreted protein levels that mediate cell-cell communication, can offer mechanistic insights without introducing perturbations. To exploit this approach, we developed the MEA-LINK platform, which utilizes simultaneous neuronal activity recordings and proteomic measurements of the supernatant of the same well (Fig. 2b). Both MEA and proteomics measurements are non-invasive, allowing longitudinal measurements across the developmental trajectory or pre-and post treatments. Using this multi-modal approach, we aimed to identify immune mediators that contribute to neuronal network activity maturation and hyperactivity control (Fig. 2b).

**Figure 2.**
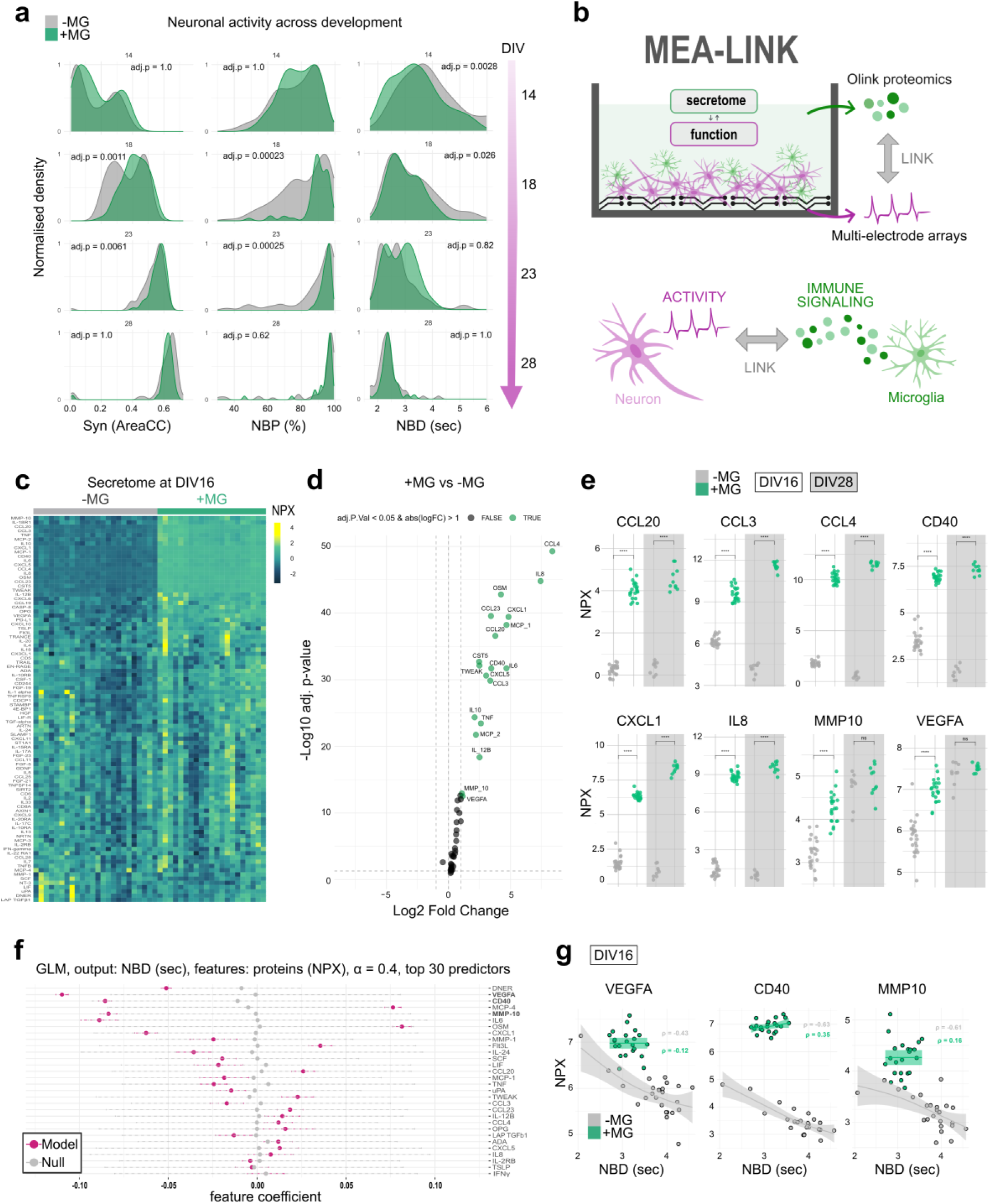
MEA-LINK leverages natural variation and identifies growth factors VEGFA, CD40, and MMP10 as predictors of network activity maturity. **(a)** Density distribution of activity metrics per condition at DIV14, DIV18, DIV23 and DIV28, n = 402 neuronal network recordings for -MG, N = 4, n = 235 neuronal network recordings for +MG, unpaired t-test with post hoc Bonferroni correction. **(b)** Schematic overview of MEA-LINK technology to identify microglia-mediated drivers of neuronal activity. **(c)** Heatmap using unsupervised clustering of normalized protein expression of all 92 Olink proteins measured in supernatant per neuronal network at DIV16, n = 24 for -MG, n = 21 for +MG. **(d)** Differential secretion analysis comparing the secretome of +MG to -MG neuronal networks at DIV16. **(e)** Normalized protein expression levels of secreted proteins at DIV16 and DIV28 per condition, n = 24 for DIV16 -MG, n = 21 for DIV16 +MG, n = 9 for DIV28 -MG, n = 10 for DIV28 +MG, unpaired t-test with post-hoc Bonferroni correction, ****p <0.0001. **(f)** Generalized linear model (GLM) of DIV16 MEA-LINK samples at alpha = 0.4 showing feature coefficients of proteins for predicting network burst duration (sec) in Ctrl group ranked by p-value. **(g)** Spearman’s correlation plots of CD40, VEGFA, and MMP10 with network burst duration at DIV16 per condition, n = 24 for -MG, n = 21 for +MG.

### Human microglia accelerate developmental increase of MMP-10 and VEGFA which predicts neuronal network burst duration

First, we analyzed the supernatant of the neuronal networks at DIV16 and DIV28 using Olink proteomics. Clustering of the 92 secreted proteins at DIV16 revealed co-variations in protein levels between wells, including a cluster of proteins that were robustly higher in +MG networks (Fig. 2c). Statistical analysis identified 19 proteins significantly elevated in +MG compared to -MG networks (Fig. 2d). Among these were the growth factors VEGFA and MMP-10, the soluble form of the receptor CD40, and, most prominently chemokines such as CL20, CCL3, CCL4, CXCL1, IL8 (Fig. 2d). We then explored how these proteins changed over time along the developmental trajectories (Fig. 2e). The levels of the chemokines and CD40 remained elevated in +MG networks at DIV28. In contrast, MMP-10 and VEGFA were no longer prominently elevated by DIV28, as the levels in -MG networks had increased to match those in +MG networks by DIV28 (Fig. 2e). We confirmed these findings using bulk transcriptomics at DIV28 and observed higher RNA expression of the chemokines and *CD40* in +MG compared to -MG networks (Suppl. Fig. 3a). As expected, *MMP10* and *VEGFA* did not show this pattern (Suppl. Fig. 3a).

We speculated that changes in secreted protein levels across development might correspond to neuronal network maturation. For a protein to serve as a meaningful maturation marker, its levels should also explain the variability in network maturation observed within a given timepoint and condition. To test this, we performed MEA-LINK at DIV16 in -MG networks to identify proteins that predict network burst duration (NBD). We focused on NBD as this MEA parameter showed substantial variability in neuronal networks early in development (Fig. 2a) and showed a robust developmental decrease in both our data and previous reports^37^. We used a generalized linear regression model on the paired activity-protein samples using eNetXplorer^45^ (Fig. 2f, Suppl. Fig. 3b). Interestingly, CD40, VEGFA, and MMP-10 were among the top predictors of NBD (Fig. 2f) and were negatively correlated with NBD, indicating that higher protein levels were associated with shorter NBDs in -MG networks (Fig. 2g). In +MG networks the elevated levels of CD40, VEGFA, and MMP-10 did not correlate with the already shortened NBD in these more mature networks, suggesting a ceiling effect (Fig. 2g).

To determine the contribution of the different cell types in the networks to the elevated levels of secreted proteins, we analyzed single-cell RNA sequencing data from +MG networks (Suppl. Fig. 3c). Chemokines and *CD40* were predominantly expressed by microglia. *VEGFA* was highest expressed in astrocytes, whereas *MMP10* was primarily expressed by a small subset of neurons. (Suppl. Fig. 3c). Taken together, microglia elevate chemokines levels in the networks that are stable throughout development and enhance secretion of CD40, neuronal MMP-10 and astrocytic VEGFA which are factors predicting neuronal network activity maturity. Interestingly, CD40, VEGFA, and MMP-10 have all been associated with neuronal growth^46–48^, which is enhanced in networks containing microglia (Fig. 1f). Thus, MEA-LINK yields biologically relevant readouts of neuron-microglia interactions and serves as a powerful tool for generating hypotheses about neuroimmune crosstalk in different contexts.

### Human microglia rescue neuronal hyperactivity in pharmacological and genetic challenges

Microglia adapt rapidly upon a change in their environment^12^ to adequately change their function to meet the requirements of this new situation. This context-dependency is important for studying microglial function^49^. Using MEA-LINK, we investigated the role of human microglia in seizure-like conditions. To induce hyperactivity, we stimulated -MG and +MG networks with kainic acid (KA) and measured neuronal network activity both during the acute phase and subsequent recovery period. Within ten minutes of KA treatment, both conditions showed robust induction of hyperactivity, characterized by rapidly repeating bursts^50^ (Fig. 3a). While we observed no difference in the acute phase of hyperactivity induction between -MG and +MG networks (Fig. 3a), the recovery period showed microglia-specific changes. Unsupervised clustering of the MFR trajectories per well showed two distinct response clusters with separation by +MG and -MG (Fig. 3b). The +MG networks showed faster recovery starting one hour post KA treatment, and reached activity levels below their initial baseline, a state that remained for at least 24 hours post hyperactivity induction (Fig. 3c). In contrast, the -MG networks recovered more slowly and returned to their baseline activity levels by 24 hours. The persistent activity reduction in the +MG networks at 24 hours was driven by a reduction in burst duration, an effect present in -MG networks (Fig. 3c). Taken together, neuronal networks containing human microglia showed enhanced recovery and long-lasting adaptive suppression of activity following pharmacologically induced hyperactivity.

**Figure 3.**
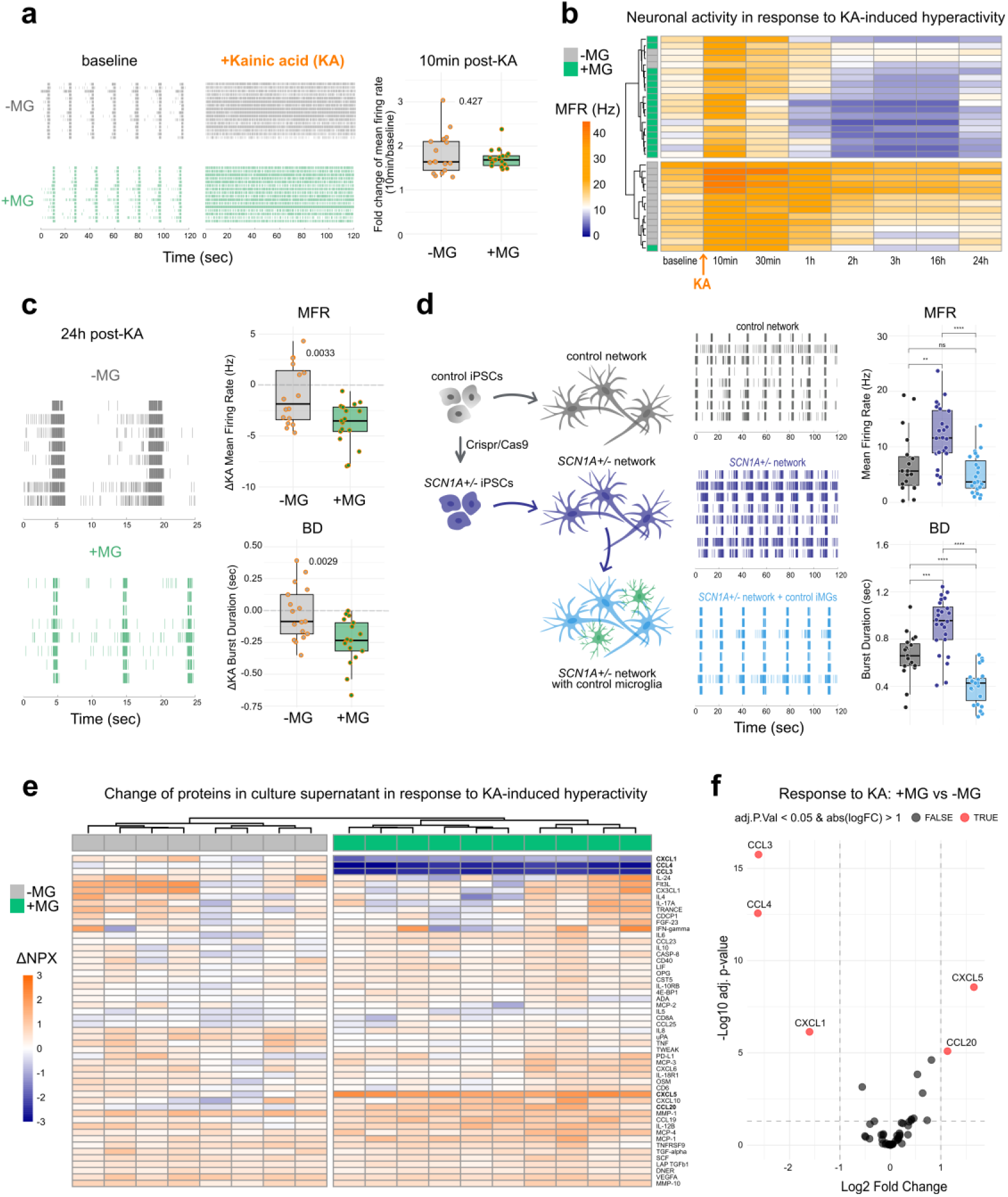
Human microglia are neuroprotective in pharmacological and genetic hyperactivity challenges. **(a)** Representative raster plots of neuronal networks at DIV28 before and after acute Kainic-acid (KA) induced hyperactivity induction (left) and quantification of fold change in mean firing rate 10 minutes post KA treatment per neuronal network, n = 16 for -MG, n = 17 for +MG, unpaired t-test (right). **(b)** Heatmap using unsupervised Euclidean clustering of neuronal networks across hyperactivity response timeline based on mean firing rate (Hz). **(c)** Representative raster plots 24 hours post KA treatment per condition (left) and quantification of the change compared to baseline in mean firing rate (MFR) and burst duration (BD) per well, n = 16 neuronal networks for -MG, n = 17 neuronal networks for +MG, **p <0.01, unpaired t-test (right). **(d)** Experimental design of experiment using *SCN1A*-deficient neuronal networks (left), representative raster plots per condition at DIV28 (middle), and quantification of mean firing rate (MFR) and burst duration (BD) at DIV28, n = 19 control networks, N = 2, n = 24 *SCN1A*+/- networks, n = 23 *SCN1A*+/- networks with control microglia, one-way ANOVA with post-hoc Bonferroni correction, **p <0.01, ***p <0.001, ****p <0.0001 (right). **(e)** Heatmap using unsupervised Euclidean clustering of neuronal networks based on the change in protein secretion (ΔNPX) 24 hours post KA treatment compared to baseline, n = 8 neuronal networks for -MG, n = 10 neuronal networks for +MG. **(f)** Differential analysis comparing protein secretion in response to KA in +MG vs -MG neuronal networks.

Next, we explored whether this neuroprotective role of microglia also extends to a genetic model of hyperactivity. We generated neuronal networks from iPSCs containing a heterozygous mutation in the *SCN1A* gene (Fig. 3d). This mutation is associated with Dravet Syndrome, a severe epileptic encephalopathy manifesting in early childhood^51^. These *SCN1A^+/-^* neuronal networks have been previously characterized as hyperactive and desynchronized^50^. By adding healthy control microglia using our co-maturation protocol^36^, we were able to rescue the increased MFR in neurons with *SCN1A* deficiency to the level of their isogenic controls (Fig. 3d). Similar to the KA challenge, the presence of microglia in the hyperactive network also led to a reduction in burst duration (Fig. 3c-d). Interestingly, the reduction in MFR was not observed in control +MG networks at baseline (Fig. 1e), highlighting the context-dependent function of microglia. In summary, we found a neuroprotective role of human microglia in both a pharmacological and genetic challenge of neuronal network hyperactivity.

To investigate how microglia-mediated protection during hyperactivity challenges is reflected at the level of the secreted protein landscape, we profiled the supernatant at baseline and 24 hours post KA administration using Olink proteomics. We calculated the change in protein secretion (ΔNPX) per well and performed unsupervised clustering analysis (Fig. 3e). The +MG networks showed a distinct proteomic response to KA and clustered separately from the -MG networks. In general, most protein levels increased after KA treatment, however CCL3, CCL4 and CXCL1 levels decreased significantly in the +MG networks (Fig. 3e). Differential analysis revealed five proteins that responded significantly different in +MG networks, and strikingly these were exclusively chemokines (Fig. 3f), suggesting a role for chemokine signaling in hyperactivity challenges.

### Microglia-dependent change in CCL4 predicts reduction in mean firing rate

To identify proteins that could potentially mediate the microglia-driven reduction in neuronal hyperactivity, we performed MEA-LINK with longitudinal sampling pre and post KA treatment, correlating changes in activity (ΔMFR) with changes in protein secretion (ΔNPX). We found that uPA, VEGFA, CCL4, and IL5 significantly predicted the MFR reduction (Fig. 4a). Among these proteins, CCL4 stood out as both microglia-dependent and showing a high fold change after KA treatment (Fig. 3f), making it a top candidate for further investigation. Indeed, correlating ΔCCL4 with ΔMFR per well revealed a positive association (Spearman’s ρ = 0.55; Fig. 4b). Notably, this correlation was exclusive to networks post hyperactivity challenge and was not observed at baseline during development (Fig. 4b), indicating that the relationship between CCL4 and neuronal activity is context-dependent.

**Figure 4.**
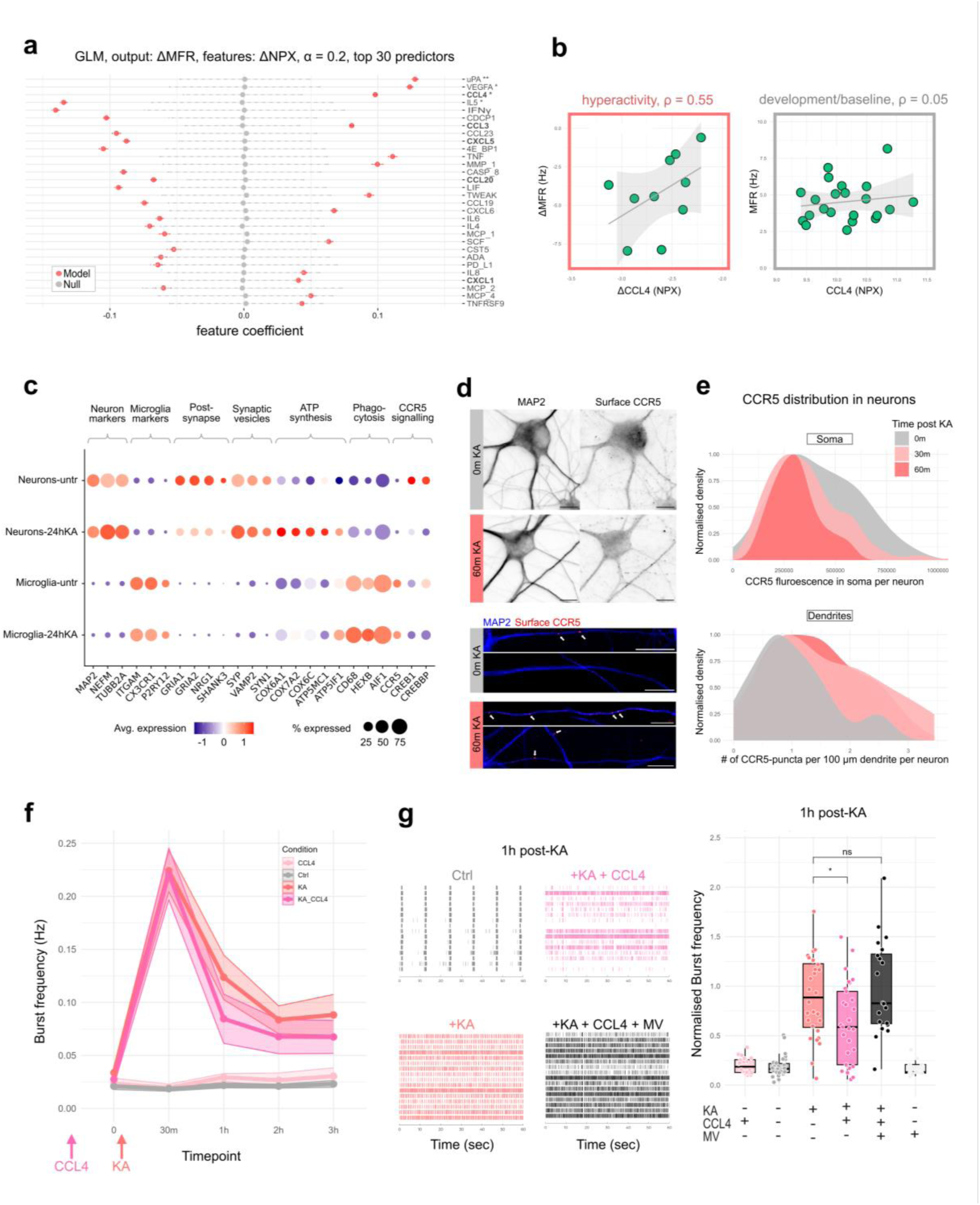
Microglial CCL4 to neuronal CCR5 signaling axis reduces neuronal activity in a hyperactivity context. **(a)** Generalized linear model (GLM) of KA-treated MEA-LINK samples at alpha = 0.2 showing feature coefficients of proteins for predicting the change in mean firing rate (Hz) ranked by p-value, microglia-dependent protein changes in bold, n = 18 paired measurements. **(b)** Spearman’s correlation of the change in CCL4 supernatant levels (Δ NPX) and the change in mean firing rate (Δ MFR) in the +MG networks after hyperactivity, n = 10 (left) and of CCL4 supernatant levels with MFR baseline, n = 22 (right). **(c)** Single-cell RNA sequencing data of cultures at baseline and 24 hours post KA treatment, N = 3, n = 49,678 single cells (45% untreated, 55% KA) showing average gene expression of selected modules in neurons and microglia untreated or KA treated. **(d)** Representative fluorescence microscope images of surface CCR5 on KA-treated neurons showing soma (top) and on dendrites (bottom), scale bars = 10 μm. **(e)** Quantification of surface CCR5 in neuronal soma (n = 46-65 neurons per timepoint) and on neuronal dendrites (n = 20 neurons per timepoint, average of 3-7 dendrites per neuron) at 0, 30, and 60 minutes after KA treatment. **(f)** Timeline of activity (burst frequency, BF) in neuronal networks post KA treatment with prior CCL4 addition (30 minutes before KA). **(g)** Representative raster plots of neuronal network activity one hour post KA treatment and additional CCL4 and CCR5-inhibitor maraviroc (MV) (left) and quantification of BF at one hour post KA treatment, N = 3, n = 34 for Ctrl, n = 20 for CCL4, n = 28 for KA, n = 29 for KA + CCL4, n = 20 for KA + CCL4 + MV, n = 10 for MV, one-way ANOVA with post-hoc Bonferroni correction, *p<0.05 (right).

Next, we focused on mechanistically understanding the role of microglial CCL4 in neuronal activity regulation. The observed reduction of CCL4 levels in the supernatant could be explained by two scenarios: less secretion of the chemokine or more uptake via the CCR5 receptor on target cells (Suppl. Fig. 3a). The strong correlation between reduction in CCL3 and CCL4 levels (Suppl. Fig. 3b), both ligands of CCR5, was hinting towards the involvement of CCR5. To explore this, we analyzed transcriptomic changes in the cultures 24 hours after KA treatment using single-cell RNA sequencing (Fig. 4c, Suppl. Fig. 3d-e). Although the mRNA levels of *CCR5* in neurons were not significantly altered at this late 24 hour timepoint, the CREB-pathway, known to be inhibited by CCR5 activation^52^, was downregulated in KA-treated neurons (Fig. 4c). Notably, the downregulation of the CREB pathway following KA stimulation was less pronounced in -MG networks (Suppl. Fig. 3f), supporting a role for microglial CCL4-CCR5 signaling in neuronal activity suppression. Functionally, neurons also showed transcriptional changes consistent with adaptive remodeling after hyperactivity: genes related to post-synaptic organization and AMPARs were downregulated, while genes involved in the synaptic vesicle cycle were upregulated (Fig. 4c). The strong upregulation in genes linked to ATP synthesis suggest higher energy demands after hyperactivity.

To examine CCR5 levels in KA-treated neurons at earlier time points, we performed immunocytochemistry for surface CCR5. Interestingly, we observed a redistribution of CCR5 from the neuronal soma towards the dendrites as early as 30 minutes after KA treatment, which became more pronounced by 60 minutes (Fig. 4d-e). These findings indicate a dynamic, activity-dependent regulation of CCR5 in human neurons.

### CCL4-CCR5 signaling reduces neuronal activity in a hyperactivity context

While MEA-LINK uncovers associations between inflammatory mediators and neuronal network activity, it does not establish causality. To directly test if the predicted CCL4 mediates activity reduction via neuronal CCR5, we designed an *in vitro* validation experiment. We postulated a context-dependent framework in which neurons upregulate CCR5 during hyperactivity^53, 54^, allowing microglial CCL4 to bind and activate a downstream cascade that reduces neuronal firing. For this model to be accurate, three conditions must be met: I) CCL4 should have no effect on activity during baseline conditions, II) CCL4 should be sufficient to reduce activity in KA-stimulated networks even in the absence of microglia, and III) blocking CCR5 should inhibit the effects of CCL4. To test these predictions, we added CCL4 to -MG networks and recorded activity after KA stimulation (Fig. 4f-g). As expected, CCL4 did not change activity at baseline, but accelerated recovery after KA-induced hyperactivity (Fig. 4f). Modelling the recovery after the hyperactivity peak with an exponential function showed a faster decay rate in the networks treated with CCL4 (Suppl. Fig. 4g). One hour post stimulation, burst frequency was significantly reduced by CCL4 addition, and this reduction was blocked by adding the competitive CCR5-inhibitor Maraviroc (MV) (Fig. 4g). Taken together, these findings demonstrate that microglial CCL4 modulates neuronal hyperactivity through neuronal CCR5 and confirmed our previously established context-dependent framework of activity control by human microglia.

## Discussion

We introduce MEA-LINK, a multi-modal platform for systematic studies of functional neuro-immune interactions in a human context. MEA-LINK is based on an optimized co-maturation model of hiPSC-derived neurons and microglia^36^. By leveraging the natural variation in biological systems, we establish correlations among protein levels and neuronal activity parameters from the same well and timepoint. The computed predictions provide insights for identifying novel targets, that can be further validated *in vitro.* Here, we applied MEA-LINK to explore microglia-neuron interactions in development and hyperactivity challenges.

### Human microglia promote neuronal network maturation

The role of microglia in neurodevelopment has been widely studied in animal models^23, 10, 15^, with a particular focus on synaptic pruning by microglia^55–57^. Recently, the necessity of synaptic pruning by microglia for healthy neurodevelopment has been challenged^58, 22^. Furthermore, it is unclear whether the findings about synaptic pruning can be generalized to all brain areas and contexts^59, 60^. Beyond synaptic refinement, studies showed that microglial support neuronal development and functions also via regulating neurogenesis^61^ or providing mitochondria^62^. We found that human microglia accelerate the development of organized neuronal network activity. Notably, there were no differences in the overall firing rate of the neurons. This emphasizes the importance of considering more sophisticated network parameters such as synchrony or network burst duration rather than just the mean firing rate in analysis of neuronal activity patterns. The faster neuronal network maturation in the presence of microglia was reflected in an accelerated increase of the growth factors VEGFA and MMP10, and levels of these proteins predicted activity maturity also in the -MG condition. These results on proteomic, morphological, and functional level point towards a neuronal network maturing role of human microglia that extends beyond the pruning of synapses.

Investigating multiple timepoints is essential when studying complex cell-cell interactions. Here, we observed marked differences in network activity around DIV18, but no differences in the same parameters by DIV28, consistent with microglia-accelerated maturation. Had only a single time point been sampled the data would have led to misleading conclusions, for example that microglia have no effect on neuronal activity. Considering this temporal complexity of microglia-neuron interaction is also relevant for rodent studies. Any difference in age of the mice studied in each experiment, can influence the results and might explain conflicting findings between different studies and laboratories.

### Human microglia rescue neuronal hyperactivity

Several studies in rodents have indicated a neuroprotective role of microglia in seizure models or epilepsy^27, 25, 24, 26^. Here, we replicate the microglial-mediated rescue of neuronal hyperactivity in a human model by employing both pharmacological stimulation and genetic representations of epilepsy. Our findings reveal that while microglia do not alter the initial induction of hyperactivity, they significantly improve the recovery phase. This is consistent with prior observations in rodent models where both the duration and severity of seizure-like activity were enhanced by microglia depletion^25^. Therefore, we hypothesize that, while microglia do not prevent seizure development, they rather have various mechanisms in place to limit the hyperactivity.

Since the presence of microglia in the neuronal networks elicited a more significant protective effect compared to addition of CCL4, it is likely that multiple mechanisms contribute to the protective function of microglia. Regulation by the identified CCL4 is one of the control pathways set in place. Next to other secreted factors, these mechanisms can also be mediated by physical cell-cell contact of microglia ramifications and synapses^63^. Using CellChat analysis^64^, we observed upregulation in complement signaling after hyperactivity (Suppl. Fig. 4h). Specifically focusing on signaling from neurons to microglia, we found that neuronal C3 to microglial CR3 (*ITGAM*) was upregulated after KA stimulation (Suppl. Fig. 4i), indicating enhanced complement-dependent remodeling after hyperactivity induction. As the complement system has been repeatedly implicated in rodent *in vivo* studies of microglia-neuron crosstalk^65, 66, 57^, this confirmed that our human iPSC-derived co-culture model reflects important aspects of microglial biology. The C3-receptor has been shown to be involved in microglia-mediated synapse elimination^65^, and removal of synapses after excessive activity might display another mechanism for activity reduction by microglia.

### Chemokines: a novel player in regulating neuronal activity

To identify potential immune modulators regulating neuronal hyperactivity, we screened 92 immune related proteins. Notably, the top hyperactivity responding proteins were all chemokines. In fact, cytokines such as IFN-γ or IL-17 have been previously implicated in neuronal activity modulation^67, 41, 68^. Correspondingly, many neurodevelopmental and psychiatric disorders have been associated with cytokine function^69, 70^. A role of chemokines in directly controlling neuronal activity is less well characterized. We identified context-dependent activity reduction via microglial CCL4 to neuronal CCR5 signaling. Our findings support a neuron-specific function of chemokines and their receptors in the brain, distinct from their established role in peripheral immune cell migration.

CCR5 has been widely studied for its role in HIV infections. In the context of the brain, it recently gained attention as a therapeutic target in stroke research^71^. Inhibiting CCR5 resulted in a protective effect against motor impairment in mice^52^. This might be partially mediated by increased neuronal activity beneficial for induction of repair mechanisms. A study exploring memory function in mice found that CCR5 signaling led to decreased excitability of neurons in acute brain slices^72^. While they link the reduced activity to a deficit in memory ensembles, the very same mechanisms might be beneficial in situations of abnormally high activity, like epilepsy. Previous studies have reported a pro-inflammatory environment, including high CCL4 levels, in epileptic brain tissue of rodents and humans^30, 29^. However, the causality remains unclear. We propose that immune activation in this context serves a homeostatic role. Chemokine signaling operates as a stress response to regulate hyperactivity rather than driving epileptogenesis. These findings suggest that simply reducing inflammation may not be an effective therapeutic approach. While excessive immune activation can be detrimental, it may occur as a consequence of system exhaustion and was intended to dampen activity in the first place. A more nuanced understanding of microglia-to-neuron signaling can lead to better-targeted interventions that balance immune response without compromising its neuroprotective role.

### Microglia-neuron signaling is context-dependent

Microglial function dynamically adapts to the brain’s needs at any given time^11^. This context-dependency should be considered in studying their role in brain function, and our findings highlight that this also holds true for microglia-neuron crosstalk. For example, CCL4 does not regulate neuronal activity under baseline conditions but plays a significant role during hyperactivity challenges. This is mediated by conditional expression and localization of CCR5. During hyperactivity challenges like stroke, NMDA or KA administration, neurons upregulate CCR5^54, 71, 53^. This mechanism ensures tight control of neuronal activity control, as it operates only in situations where two conditions are met: microglial secretion of CCL4 and neuronal upregulation of CCR5 at the level of the dendrites. Our results emphasize the importance of studying immune functions in the brain beyond baseline conditions. Many immune regulatory mechanisms may remain inactive until they are needed, making it essential to investigate their roles under various physiological and pathological challenges. While epilepsy serves as an example in this study, the principle of context dependency applies broadly to other neurological conditions.

### Limitations and future directions

In this study, we screened for soluble immune modulators of activity. However, microglia-driven regulation of network activity can also occur at the level of synapse remodeling, uptake of debris, metabolic support or other physical cell-cell interactions^57, 56, 63, 62, 73^. For initial proof-of-concept of the MEA-LINK platform and the establishment of relevant correlations among parameters, we used the 92-taget Olink panel. However, in the future expanding to a bigger discovery panel will offer capturing a broader view of neuro-immune interactions and how soluble factors shape neuronal activity. Furthermore, the hiPSC-derived co-culture model used in this study did not include GABAergic neurons. Recently, microglia have been shown to specifically interact with inhibitory synapses and thus shape inhibitory circuits^74, 61, 75^. Therefore, exploiting the possibility to also include hiPSC-derived inhibitory neurons^76^ in our microglia-neuron co-culture model will allow for a more complete understanding of neuronal circuit control by microglia in health and disease. Despite these limitations, this study introduces a new platform for systematically screening functional neuro-immune interactions in a human context. We used MEA-LINK to understand microglia-mediated network control in development and hyperactivity challenges. Importantly, this multi-modal time-series approach can be applied beyond epilepsy research. Various perturbations such as patient-derived cerebrospinal fluid, immune-related stimulations, or protein aggregates like amyloid-beta/a-synuclein can be screened with MEA-LINK to explore neuro-immune interactions in different CNS disorders.

## Methods

### Human induced pluripotent stem cell derived neuron-microglia co-cultures

To generate neuronal networks with microglia derived from human induced pluripotent stem cells (hiPSCs), we used a previously published protocol^36^. A detailed characterization and step-by-step explanation of the optimized protocol including troubleshooting tips can be found there^36^. In brief, neuronal differentiation was induced by doxycycline-dependent overexpression of *Neurogenin*-2 (*Ngn2*). On MEA plates, droplet-plating was performed to ensure successful coverage of the electrodes. HiPSC-derived pre-microglia were added in a 1:5 ratio to the neurons after doxycycline wash out at DIV9. Then, both cell types co-matured together until DIV28 in a newly formulated medium compatible with differentiation of both cell types.

The healthy control hiPSC line used in this study was GM25256, obtained from the Coriell Institute (RRID: CVCL_Y803). This line is fibroblast-derived from clinically normal 30 year old male. The *SCN1A*-deficient hiPSC line was generated with Crispr/Cas9 from GM25256. Full characterization and information regarding the *SCN1A* mutation can be found in a previous publication^50^.

### Micro-electrode array recording and analysis

To record neuronal network activity, cells were plated on CytoView 48-well or 96-well micro-electrode arrays (MEAs) and recorded using the Axion Maestro Pro MEA system (Axion Biosystems). Before measurements all plates acclimatized for a minimum of 10 minutes to ensure a stable temperature of 37°C and 5% CO_2_. Recordings of 5 min were always performed ∼24 hours after medium change as this procedure influences neuronal firing. All corresponding supernatants for Olink proteomics were taken less than 5 minutes after recordings for an exact time match. Not more than 50 µl of supernatant was taken from the MEA wells (300 µl total) to not disturb activity dynamics during an experiment. In the MEA recording spikes were detected when voltage deflections exceeded an adaptive threshold of ±6 standard deviations from the noise level. Processing of the MEA data was done using the Axion Biosystems NeuralMetric Tool. Bursts were detected when at least 5 consecutive spikes had a maximum inter-spike interval of 100 ms each. The envelope network burst detection algorithm of was used with these settings: threshold = 1.5, minimum inter burst interval = 100 ms, minimum number of active electrodes = 70%, burst inclusion = 75%.

Neuronal networks with <1 Hz mean firing rate (MFR) or <1 network bursts (NBs) within 5 minutes of recording were excluded from analysis. Filtered wells were ∼5% of total recordings and we continued with 1,137 network recordings for downstream analysis. Since many output MEA parameters are derived from similar metrics and thus highly correlated, we explored parameter-parameter correlations (Suppl. Fig. 1c-d) and decided to include these parameters: MFR, Burst duration (BD), Burst frequency (BF), Network burst duration (NBD), Network burst percentage (NBP) and Area under normalized cross correlation as Synchrony measure (Syn). We did not observe an effect of the location of the well (border or not) within the plate on the network activity parameters (Suppl. Fig. 1e). Heatmaps per well across time were plotted using the *pheatmap* R package and Euclidean clustering. Since principal component-cofactor correlations and variance partition analysis revealed that neuronal batch was a source of variation (Suppl. Fig. 1e) we corrected for neuronal batch in the developmental heatmap to minimize batch-related clustering. In the CCL4 validation experiments, we normalized each of the three batches for the magnitude in response to KA (Fig. 4g). All other plots show non-normalized values on the true scale of the activity metric.

### Pharmacology

All reagents were prepared into stocks and stored at −20°C. We chose Kainic acid to induce a hyperactivity response in the neuronal network at DIV28 (KA, 5 µM, Sigma 58002-62-3). For the *in vitro* validation experiments, Maraviroc was added 30 minutes before KA treatment (1 μM, Selleckchen, S2003) and CCL4 (100 ng/ml, Peprotech, 300-09) was added 5 minutes before KA treatment. The working dilution was added directly to the wells 5 minutes prior to the MEA recordings under sterile conditions. For inhibition of glutamate receptors, the MEA wells were treated with D-2-amino-5-phosphonovalerate (D-AP5, 100 μM, Tocris, 0106), then 1-naphthyl acetyl spermine trihydrochloride (Naspm, 10 μM, Tocris, 2766) and finally 2,3-dioxo-6-nitro-1,2,3,4-tetrahydrobenzo[F]quinoxaline-7-sulfonamide (NBQX, 50 μM, Tocris, 0373). If possible and recommended by the manufacturer, compounds were diluted in ultra-pure water. Otherwise DMSO was used. For all pharmacological treatments and experiments, the amount of DMSO in the cell culture medium was ≤0.5% v/v.

### Immunocytochemistry

Unless stated otherwise, subsequent steps were performed at room temperature and solutions were made in 1XPBS. Cultured cells were fixed by using a gradient of 2% and subsequently 4% paraformaldehyde (PFA) supplemented with 4% sucrose for 15 minutes in total. Then, cells were washed three times with 1XPBS, and permeabilized with 0.2% Triton (Sigma-Aldrich, T8787) for 10 minutes. To block nonspecific binding sites, cells were incubated with 3% bovine serum albumin (BSA, Sigma, A7906) for 1h. Primary antibodies were diluted in the same buffer and incubated overnight at 4°C. The following primary antibodies and dilutions were used: guinea pig anti-MAP2 (1:1000, Synaptic Systems, 188004), mouse anti-panAxonal (1:1000, BioLegend, 837904), rabbit anti-GFAP (1:1000; Sigma-Aldrich, AB5804), rabbit anti-Synapsin I (1:500, Sigma-Aldrich, AB1543P), mouse anti-Homer1b/c (1:200, Synaptic Systems, 160111), chicken anti-NeuN (1:500, Sigma-Aldrich, ABN91), mouse anti-CCR5 (CD195, 1:100, Invitrogen, 14-1957-82).

Secondary antibodies conjugated to either Alexa 488, Alexa 568 or Alexa 647 (Invitrogen) were incubated for 1h at 1:1000 in 1% BSA. Cell nuclei were stained with Hoechst (0.01%, ThermoFisher, H3570) for 10 minutes. After three final washes, the coverslips were mounted in DAKO (Agilent, S3023) on microscope slides and imaged on a Zeiss Axio Imager Z2.

### Somatodendritic reconstructions and morphometrical analyses

To analyze neuronal morphology, we imaged MAP2-stained neurons across DIV12, DIV14, DIV21, and DV28 at x20 magnification. We excluded neurons that were completely isolated from other neurons at the edge of the coverslip, as well as neurons that had overlapping soma with neighboring neurons to allow for confident tracing of the primary dendrites. Somatodendritic reconstructions were performed using a vector based morphometrical reconstruction software (NeuroLucida 360, Version 11, MBF-Bioscience, Williston, ND, USA). The reconstruction was performed blinded and only on neurons that could be confidently separated from their neighboring neurons. R was used for data preprocessing of the NeuroLucida output and subsequent analysis. We quantified the number of primary dendrites, average length per dendrite, total dendritic length per neuron and the number of dendritic nodes per neuron. To investigate the complexity of dendrites in regard to their distance from the soma, we performed Sholl analysis and quantified the number of intersections per 10 µm interval.

### Synapse quantification

To quantify the number of synapses, the cultures were stained for MAP2, Synapsin1, Homer1 at DIV17, 21, and 28. Images were taken at x63 magnification using oil, and with apotome settings to reduce background noise. The number of (co-localized) puncta were quantified using an adapted version of the SynBot plug-in^77^. The adaptation was made to quantify synapses per µm dendrite using ROIs based on the MAP2 staining. The noise reduction parameters were adjusted to a Gaussian Blur Sigma of 0.57 and a Rolling Ball Background Subtraction Radius of 50. Ilastik-based thresholding training was performed per DIV using 3 images of both conditions (-MG and +MG). Conditions were analyzed using the same training per DIV to ensure comparability between conditions. Co-localization was determined by pixel overlap.

### CCR5 quantification

We analyzed the distribution of CCR5 after Kainic acid (KA) treatment in neurons at DIV28. The cultures were treated with KA (5 µM, Sigma 58002-62-3) for 0, 30, or 60 minutes before fixation. The neurons were stained for CCR5 before permeabilizing to ensure surface staining. After washing and subsequent permeabilization, the neurons were further stained with MAP2, NeuN, and Hoechst. For analysis of the soma, images were taken at x20 magnification. The soma borders were defined using the NeuN staining. For analysis of the dendrites per dendritic length, images were taken at x63 magnification using oil, and with apotome settings to reduce background noise. Only the larger bright CCR5 puncta were quantified (see arrow annotation in Fig. 4d) to avoid ambiguity in the analysis. Per neuron, an average of 3-7 dendrites was taken.

### Olink proteomics

To assess the secretome of the cultures, we performed Olink® proteomics on the supernatants using the Target 96 inflammation panel I. This Proximity Extension Assay (PEA) based technology uses antibody pairs with oligonucleotide-labels, which are amplified by real-time quantitative PCR. Using this approach allowed for sampling of low volumes, which was essential for our time-series samples of the same well. Medium changes and stimulations of microglia were always performed 24 hours before sampling. For the MEA-LINK measurements, supernatant was sampled less than 5 minutes after the MEA-recording. All samples were stored at −20°C until further processing. The Olink® assay was performed according to the manufacturer’s instructions and the data was normalized using the internal controls on the same plate. For the secretome at DIV16 we excluded proteins with a missing frequency above 97% (2 out of 96), meaning that at least one out of the 48 samples need to show abundance of a given protein. The protein levels are shown in relative protein expression value in normalized protein expression (NPX) units. We further normalized the proteins using a cell-free medium sample for each respective cell type and experiment. For calculating ΔNPX after KA stimulation, proteins with a baseline NPX below 1.5 in all samples were excluded. For data analysis, heatmaps were generated using unsupervised Euclidean clustering with the *pheatmap* R package.

### RNA-sequencing

MEA wells were washed with 1xPBS and collected using DNA/RNA shield (Zymo 20 Research; ZY-R1200-125). RNA was isolated using the Quick-RNA Microprep kit (Zymo 20 Research, ZY-R1051) according to manufacturer’s instructions. NEBNext® Ultra™ II RNA Library Prep Kit (Illumina) was used for library preparation as per manufacturer’s recommendations. Sequencing was performed on a NovaSeq X+ (Illumina) at 2×150bp configuration with coverage of 20 million reads per sample. Raw RNA-sequencing reads were processed using the nf-core/rnaseq pipeline (v3.18) implemented in Nextflow^78^. Quality control was performed using FastQC and MultiQC and adapter trimming was conducted with Trim Galore. Reads were aligned to the human reference genome (GRCh38) using STAR with default parameters. The transcript levels were quantified with Salmon. Log2 transformation with a pseudo-count of +1 was performed on counts per million (CPM) to achieve normal distribution. Heatmaps were generated using Euclidean clustering in the *pheatmap* R package.

### Single cell RNA-sequencing

At DIV28, the neuron-microglia co-cultures were dissociated using Accutase (20 minutes at 37°C). We performed the KA stimulation 24 hours before harvesting the samples to streamline the harvesting for untreated and treated wells. Two to three technical replicates were used per condition. The single-cell suspensions were processed using the Chromium Next GEM Single Cell Fixed RNA Sample Preparation Kit (10x Genomics, PN-1000414) according to the manufacturer’s guidelines. The samples were fixed for 24 hours at 4°C and stored with Enhancer at −80° until further processing. We prepared the libraries using the Chromium Single Cell Flex Kit (10x Genomics) following the manufacturer’s protocol (CG000527 Rev F). Raw sequencing data were processed and de-multiplexed using the Cell Ranger pipeline (10x Genomics, Cellranger v. 9.0.0) with human reference genome refdata-gex-GRCh38-2024-A and Chromium_Human_Transcriptome_Probe_Set_v1.1.0_GRCh38-2024-A. Downstream analyses, including quality control, normalization, library integration, doublet detection, clustering, and differential gene expression analysis, were performed using Seurat (v4.3.0) in R. Quality control criteria included the removal of cells with fewer than 200 genes detected, fewer than 500 UMIs or with more than 10% mitochondrial gene content. To remove doublets, cells with an UMI count two standard deviations above the mean were excluded. Doublets were identified by their characteristic floating small cluster shape and mixed gene expression profiles. Harmony^79^ (v1.2.3) was used for integration of the three batches/libraries. Cell types were annotated based on the top differentially expressed genes of the Seurat clusters. To predict ligand-receptor pairs based on the single cell transcriptomes and infer cell-cell communication CellChat (v2.1.2) was used^64^.

### Elastic net of generalized linear models

The predictions of neuronal network activity by the Olink measured proteins were conducted using the eNetXplorer R package^45^. NPX and activity values were matched by their Well ID. The best α value (0 = ridge, 1 = lasso) for the generalized linear model (GLM) was determined by model accuracy as quality function (Suppl. Fig. 3b, 4c). The predictors for the corresponding α were ordered according to their p-value and the feature coefficient was plotted in a caterpillar plot comparing the model to the null reference calculated without predictor variables.

### Statistical analysis

Statistical analyses were performed with R (RStudio, R version 4.2.0) or GraphPad PRISM 8.0.0 (GraphPad Software, Inc., CA, USA). All values are reported as mean ± standard error of the mean (SEM), unless stated otherwise. In all figures, *P*-values are indicated as follows: \**P*<0.05, ** *P* <0.005, \*\*\**P* <0.0005, **** *P* <0.0001. For comparisons between two conditions at one time point, an unpaired t-test was performed. For comparison of two or more conditions with repeated measures, we used repeated measures two-way ANOVA. Any time multiple conditions or time points were compared, we applied post-hoc Bonferroni correction.

## Data Availability

The source data for all figures in the main text (Figures 1–4) are available in Supplementary table 1. All raw datasets generated during the current study are available from the corresponding author on reasonable request.

## Acknowledgments

We thank all members of the de Witte, Nadif Kasri, and Tsang labs for their valuable input during discussions. This work was supported by the Hypatia Grant from the RadboudUMC (L.D.W), the Simons Foundation #00010410 (L.D.W), the SFARI grant #890042 (N.N.K.), the Stichting de Drie Lichten (A.M) and the Dutch Research Council (NWO, ZonMW Open Competitie #09120012110034 and #OCENW.XS24.2.134, L.D.W).

## References

1. Hubel, D.H., and Wiesel, T.N. (1959). Receptive fields of single neurons in the cat’s striate cortex. The Journal of physiology 148, 574–591. 10.1113/jphysiol.1959.sp006308.

2. Siskind, G.W., and Benacerraf, B. (1969). Cell selection by antigen in the immune response. Advances in immunology 10, 1–50. 10.1016/s0065-2776(08)60414-9.

3. Nabavi, S., Fox, R., Proulx, C.D., Lin, J.Y., Tsien, R.Y., and Malinow, R. (2014). Engineering a memory with LTD and LTP. Nature 511, 348–352. 10.1038/nature13294.

4. Lam, N., Lee, Y., and Farber, D.L. (2024). A guide to adaptive immune memory. Nature reviews. Immunology 24, 810–829. 10.1038/s41577-024-01040-6.

5. Renart, A., Song, P., and Wang, X.-J. (2003). Robust spatial working memory through homeostatic synaptic scaling in heterogeneous cortical networks. Neuron 38, 473–485. 10.1016/S0896-6273(03)00255-1.

6. van Parijs, L., and Abbas, A.K. (1998). Homeostasis and self-tolerance in the immune system: turning lymphocytes off. Science (New York, N.Y.) 280, 243–248. 10.1126/science.280.5361.243.

7. Cannon, W.B. (1929). Organization for physiological homeostasis. Physiological Reviews 9, 399–431. 10.1152/physrev.1929.9.3.399.

8. Nayak, D., Roth, T.L., and McGavern, D.B. (2014). Microglia development and function. Annual review of immunology 32, 367–402. 10.1146/annurev-immunol-032713-120240.

9. Borst, K., Dumas, A.A., and Prinz, M. (2021). Microglia: Immune and non-immune functions. Immunity 54, 2194–2208. 10.1016/j.immuni.2021.09.014.

10. Prinz, M., Jung, S., and Priller, J. (2019). Microglia Biology: One Century of Evolving Concepts. Cell 179, 292–311. 10.1016/j.cell.2019.08.053.

11. Matcovitch-Natan, O., Winter, D.R., Giladi, A., Vargas Aguilar, S., Spinrad, A., Sarrazin, S., Ben-Yehuda, H., David, E., Zelada González, F., and Perrin, P., et al. (2016). Microglia development follows a stepwise program to regulate brain homeostasis. Science (New York, N.Y.) 353, aad8670. 10.1126/science.aad8670.

12. Gosselin, D., Skola, D., Coufal, N.G., Holtman, I.R., Schlachetzki, J.C.M., Sajti, E., Jaeger, B.N., O’Connor, C., Fitzpatrick, C., and Pasillas, M.P., et al. (2017). An environment-dependent transcriptional network specifies human microglia identity. Science (New York, N.Y.) 356. 10.1126/science.aal3222.

13. Umpierre, A.D., and Wu, L.-J. (2021). How microglia sense and regulate neuronal activity. Glia 69, 1637–1653. 10.1002/glia.23961.

14. Priller, J., and Prinz, M. (2019). Targeting microglia in brain disorders. Science (New York, N.Y.) 365, 32–33. 10.1126/science.aau9100.

15. Salter, M.W., and Stevens, B. (2017). Microglia emerge as central players in brain disease. Nat Med 23, 1018–1027. 10.1038/nm.4397.

16. Colonna, M., and Butovsky, O. (2017). Microglia Function in the Central Nervous System During Health and Neurodegeneration. Annual review of immunology 35, 441–468. 10.1146/annurev-immunol-051116-052358.

17. Paolicelli, R.C., Sierra, A., Stevens, B., Tremblay, M.-E., Aguzzi, A., Ajami, B., Amit, I., Audinat, E., Bechmann, I., and Bennett, M., et al. (2022). Microglia states and nomenclature: A field at its crossroads. Neuron 110, 3458–3483. 10.1016/j.neuron.2022.10.020.

18. Villani, A.-C., Sarkizova, S., and Hacohen, N. (2018). Systems Immunology: Learning the Rules of the Immune System. Annual review of immunology 36, 813–842. 10.1146/annurev-immunol-042617-053035.

19. Mulè, M.P., Martins, A.J., Cheung, F., Farmer, R., Sellers, B.A., Quiel, J.A., Jain, A., Kotliarov, Y., Bansal, N., and Chen, J., et al. (2024). Integrating population and single-cell variations in vaccine responses identifies a naturally adjuvanted human immune setpoint. Immunity 57, 1160–1176.e7. 10.1016/j.immuni.2024.04.009.

20. Eling, N., Morgan, M.D., and Marioni, J.C. (2019). Challenges in measuring and understanding biological noise. Nature reviews. Genetics 20, 536–548. 10.1038/s41576-019-0130-6.

21. Tsang, J.S. (2015). Utilizing population variation, vaccination, and systems biology to study human immunology. Trends in immunology 36, 479–493. 10.1016/j.it.2015.06.005.

22. O’Keeffe, M., Booker, S.A., Walsh, D., Li, M., Henley, C., Simões de Oliveira, L., Liu, M., Wang, X., Banqueri, M., and Ridley, K., et al. (2025). Typical development of synaptic and neuronal properties can proceed without microglia in the cortex and thalamus. Nature neuroscience 28, 268–279. 10.1038/s41593-024-01833-x.

23. Zengeler, K.E., and Lukens, J.R. (2021). Innate immunity at the crossroads of healthy brain maturation and neurodevelopmental disorders. Nature reviews. Immunology 21, 454–468. 10.1038/s41577-020-00487-7.

24. Eyo, U.B., Murugan, M., and Wu, L.-J. (2017). Microglia-Neuron Communication in Epilepsy. Glia 65, 5–18. 10.1002/glia.23006.

25. Gibbs-Shelton, S., Benderoth, J., Gaykema, R.P., Straub, J., Okojie, K.A., Uweru, J.O., Lentferink, D.H., Rajbanshi, B., Cowan, M.N., and Patel, B., et al. (2023). Microglia play beneficial roles in multiple experimental seizure models. Glia 71, 1699–1714. 10.1002/glia.24364.

26. Liu, M., Jiang, L., Wen, M., Ke, Y., Tong, X., Huang, W., and Chen, R. (2020). Microglia depletion exacerbates acute seizures and hippocampal neuronal degeneration in mouse models of epilepsy. American journal of physiology. Cell physiology 319, C605–C610. 10.1152/ajpcell.00205.2020.

27. Badimon, A., Strasburger, H.J., Ayata, P., Chen, X., Nair, A., Ikegami, A., Hwang, P., Chan, A.T., Graves, S.M., and Uweru, J.O., et al. (2020). Negative feedback control of neuronal activity by microglia. Nature 586, 417–423. 10.1038/s41586-020-2777-8.

28. Lloyd, A.F., Martinez-Muriana, A., Davis, E., Daniels, M.J.D., Hou, P., Mancuso, R., Brenes, A.J., Sinclair, L.V., Geric, I., and an Snellinx, et al. (2024). Deep proteomic analysis of microglia reveals fundamental biological differences between model systems. Cell reports 43, 114908. 10.1016/j.celrep.2024.114908.

29. Liu, Q., Shen, C., Dai, Y., Tang, T., Hou, C., Yang, H., Wang, Y., Xu, J., Lu, Y., and Wang, Y., et al. (2024). Single-cell, single-nucleus and xenium-based spatial transcriptomics analyses reveal inflammatory activation and altered cell interactions in the hippocampus in mice with temporal lobe epilepsy. Biomarker research 12, 103. 10.1186/s40364-024-00636-3.

30. Kumar, P., Lim, A., Hazirah, S.N., Chua, C.J.H., Ngoh, A., Poh, S.L., Yeo, T.H., Lim, J., Ling, S., and Sutamam, N.B., et al. (2022). Single-cell transcriptomics and surface epitope detection in human brain epileptic lesions identifies pro-inflammatory signaling. Nature neuroscience 25, 956–966. 10.1038/s41593-022-01095-5.

31. Hanger, B., Couch, A., Rajendran, L., Srivastava, D.P., and Vernon, A.C. (2020). Emerging Developments in Human Induced Pluripotent Stem Cell-Derived Microglia: Implications for Modelling Psychiatric Disorders With a Neurodevelopmental Origin. Frontiers in psychiatry 11, 789. 10.3389/fpsyt.2020.00789.

32. Zhang, Y., Pak, C., Han, Y., Ahlenius, H., Zhang, Z., Chanda, S., Marro, S., Patzke, C., Acuna, C., and Covy, J., et al. (2013). Rapid single-step induction of functional neurons from human pluripotent stem cells. Neuron 78, 785–798. 10.1016/j.neuron.2013.05.029.

33. Dräger, N.M., Sattler, S.M., Huang, C.T.-L., Teter, O.M., Leng, K., Hashemi, S.H., Hong, J., Aviles, G., Clelland, C.D., and Zhan, L., et al. (2022). A CRISPRi/a platform in human iPSC-derived microglia uncovers regulators of disease states. Nature neuroscience 25, 1149–1162. 10.1038/s41593-022-01131-4.

34. Jäntti, H., Kistemaker, L., Buonfiglioli, A., Witte, L.D. de, Malm, T., and Hol, E.M. (2024). Emerging Models to Study Human Microglia In vitro. Advances in neurobiology 37, 545–568. 10.1007/978-3-031-55529-9_30.

35. McQuade, A., Coburn, M., Tu, C.H., Hasselmann, J., Davtyan, H., and Blurton-Jones, M. (2018). Development and validation of a simplified method to generate human microglia from pluripotent stem cells. Molecular neurodegeneration 13, 67. 10.1186/s13024-018-0297-x.

36. Mordelt, A., Schuurmans, I.M.E., Scheefhals, N., Hommersom, M.P., Slottje, K., Mast, K., Graziani, M., Gonzalez, C.O., Wingens, L.J.A., and Schubert, D., et al. (2025). Long-term co-maturation of stem cell-derived microglia and neuronal networks: an optimized platform to assess human microglial contribution to neuronal function. BioRxiv, 10.1101/2025.10.24.684345.

37. Mossink, B., Verboven, A.H.A., van Hugte, E.J.H., Klein Gunnewiek, T.M., Parodi, G., Linda, K., Schoenmaker, C., Kleefstra, T., Kozicz, T., and van Bokhoven, H., et al. (2021). Human neuronal networks on micro-electrode arrays are a highly robust tool to study disease-specific genotype-phenotype correlations in vitro. Stem cell reports 16, 2182–2196. 10.1016/j.stemcr.2021.07.001.

38. Ling, E., Nemesh, J., Goldman, M., Kamitaki, N., Reed, N., Handsaker, R.E., Genovese, G., Vogelgsang, J.S., Gerges, S., and Kashin, S., et al. (2024). A concerted neuron-astrocyte program declines in ageing and schizophrenia. Nature 627, 604–611. 10.1038/s41586-024-07109-5.

39. Hommersom, M.P., Doorn, N., Puvogel, S., Lewerissa, E.I., Mordelt, A., Ciptasari, U., Kampshoff, F., Dillen, L., van Beusekom, E., and Oudakker, A., et al. (2025). CACNA1A haploinsufficiency leads to reduced synaptic function and increased intrinsic excitability. Brain : a journal of neurology 148, 1286–1301. 10.1093/brain/awae330.

40. Turner, M.D., Nedjai, B., Hurst, T., and Pennington, D.J. (2014). Cytokines and chemokines: At the crossroads of cell signalling and inflammatory disease. Biochimica et biophysica acta 1843, 2563–2582. 10.1016/j.bbamcr.2014.05.014.

41. Bellingacci, L., Canonichesi, J., Mancini, A., Parnetti, L., and Di Filippo, M. (2023). Cytokines, synaptic plasticity and network dynamics: a matter of balance. Neural regeneration research 18, 2569–2572. 10.4103/1673-5374.371344.

42. Sowa, J.E., and Tokarski, K. (2021). Cellular, synaptic, and network effects of chemokines in the central nervous system and their implications to behavior. Pharmacological reports : PR 73, 1595–1625. 10.1007/s43440-021-00323-2.

43. Demmings, M.D., Da Silva Chagas, L., Traetta, M.E., Rodrigues, R.S., Acutain, M.F., Barykin, E., Datusalia, A.K., German-Castelan, L., Mattera, V.S., and Mazengenya, P., et al. (2025). (Re)building the nervous system: A review of neuron-glia interactions from development to disease. Journal of neurochemistry 169, e16258. 10.1111/jnc.16258.

44. Cabrera-Garcia, D., Warm, D., La Fuente, P. de, Fernández-Sánchez, M.T., Novelli, A., and Villanueva-Balsera, J.M. (2021). Early prediction of developing spontaneous activity in cultured neuronal networks. Scientific reports 11, 20407. 10.1038/s41598-021-99538-9.

45. Candia, J., and Tsang, J.S. (2019). eNetXplorer: an R package for the quantitative exploration of elastic net families for generalized linear models. BMC bioinformatics 20, 189. 10.1186/s12859-019-2778-5.

46. Mackenzie, F., and Ruhrberg, C. (2012). Diverse roles for VEGF-A in the nervous system. Development (Cambridge, England) 139, 1371–1380. 10.1242/dev.072348.

47. Fujioka, H., Dairyo, Y., Yasunaga, K.-I., and Emoto, K. (2012). Neural functions of matrix metalloproteinases: plasticity, neurogenesis, and disease. Biochemistry research international 2012, 789083. 10.1155/2012/789083.

48. Chen, L., Maures, T.J., Jin, H., Huo, J.S., Rabbani, S.A., Schwartz, J., and Carter-Su, C. (2008). SH2B1beta (SH2-Bbeta) enhances expression of a subset of nerve growth factor-regulated genes important for neuronal differentiation including genes encoding urokinase plasminogen activator receptor and matrix metalloproteinase 3/10. Molecular endocrinology (Baltimore, Md.) 22, 454–476. 10.1210/me.2007-0384.

49. Depp, C., Doman, J.L., Hingerl, M., Xia, J., and Stevens, B. (2025). Microglia transcriptional states and their functional significance: Context drives diversity. Immunity 58, 1052–1067. 10.1016/j.immuni.2025.04.009.

50. van Hugte, E.J.H., Lewerissa, E.I., Wu, K.M., Scheefhals, N., Parodi, G., van Voorst, T.W., Puvogel, S., Kogo, N., Keller, J.M., and Frega, M., et al. (2023). SCN1A-deficient excitatory neuronal networks display mutation-specific phenotypes. Brain : a journal of neurology 146, 5153–5167. 10.1093/brain/awad245.

51. Deprez, L., an Jansen, and Jonghe, P. de (2009). Genetics of epilepsy syndromes starting in the first year of life. Neurology 72, 273–281. 10.1212/01.wnl.0000339494.76377.d6.

52. Wu, Q.-L., Cui, L.-Y., Ma, W.-Y., Wang, S.-S., Zhang, Z., Feng, Z.-P., Sun, H.-S., Chu, S.-F., He, W.-B., and Chen, N.-H. (2023). A novel small-molecular CCR5 antagonist promotes neural repair after stroke. Acta pharmacologica Sinica 44, 1935–1947. 10.1038/s41401-023-01100-y.

53. Mennicken, F., Chabot, J.-G., and Quirion, R. (2002). Systemic administration of kainic acid in adult rat stimulates expression of the chemokine receptor CCR5 in the forebrain. Glia 37, 124–138. 10.1002/glia.10021.

54. Galasso, J.M., Harrison, J.K., and Silverstein, F.S. (1998). Excitotoxic brain injury stimulates expression of the chemokine receptor CCR5 in neonatal rats. The American journal of pathology 153, 1631–1640. 10.1016/S0002-9440(10)65752-5.

55. Faust, T.E., Gunner, G., and Schafer, D.P. (2021). Mechanisms governing activity-dependent synaptic pruning in the developing mammalian CNS. Nature reviews. Neuroscience 22, 657–673. 10.1038/s41583-021-00507-y.

56. Paolicelli, R.C., Bolasco, G., Pagani, F., Maggi, L., Scianni, M., Panzanelli, P., Giustetto, M., Ferreira, T.A., Guiducci, E., and Dumas, L., et al. (2011). Synaptic pruning by microglia is necessary for normal brain development. Science (New York, N.Y.) 333, 1456–1458. 10.1126/science.1202529.

57. Schafer, D.P., Lehrman, E.K., Kautzman, A.G., Koyama, R., Mardinly, A.R., Yamasaki, R., Ransohoff, R.M., Greenberg, M.E., Barres, B.A., and Stevens, B. (2012). Microglia sculpt postnatal neural circuits in an activity and complement-dependent manner. Neuron 74, 691–705. 10.1016/j.neuron.2012.03.026.

58. Surala, M., Soso-Zdravkovic, L., Munro, D., Rifat, A., Ouk, K., Vida, I., Priller, J., and Madry, C. (2024). Lifelong absence of microglia alters hippocampal glutamatergic networks but not synapse and spine density. EMBO reports 25, 2348–2374. 10.1038/s44319-024-00130-9.

59. Pereira-Iglesias, M., Maldonado-Teixido, J., Melero, A., Piriz, J., Galea, E., Ransohoff, R.M., and Sierra, A. (2025). Microglia as hunters or gatherers of brain synapses. Nature neuroscience 28, 15–23. 10.1038/s41593-024-01818-w.

60. Mordelt, A., and Witte, L.D. de (2023). Microglia-mediated synaptic pruning as a key deficit in neurodevelopmental disorders: Hype or hope? Current Opinion in Neurobiology 79, 102674. 10.1016/j.conb.2022.102674.

61. Yu, D., Jain, S., Wangzhou, A., Zhu, B., Shao, W., Coley-O’Rourke, E.J., Florencio, S. de, Kim, J., Choi, J.J.-Y., and Paredes, M.F., et al. (2025). Microglia regulate GABAergic neurogenesis in prenatal human brain through IGF1. Nature. 10.1038/s41586-025-09362-8.

62. Scheiblich, H., Eikens, F., Wischhof, L., Opitz, S., Jüngling, K., Cserép, C., Schmidt, S.V., Lambertz, J., Bellande, T., and Pósfai, B., et al. (2024). Microglia rescue neurons from aggregate-induced neuronal dysfunction and death through tunneling nanotubes. Neuron 112, 3106–3125.e8. 10.1016/j.neuron.2024.06.029.

63. Eyo, U.B., Haruwaka, K., Mo, M., Campos-Salazar, A.B., Wang, L., Speros, X.S., Sabu, S., Xu, P., and Wu, L.-J. (2021). Microglia provide structural resolution to injured dendrites after severe seizures. Cell reports 35, 109080. 10.1016/j.celrep.2021.109080.

64. Jin, S., Plikus, M.V., and Nie, Q. (2025). CellChat for systematic analysis of cell-cell communication from single-cell transcriptomics. Nature protocols 20, 180–219. 10.1038/s41596-024-01045-4.

65. Anderson, S.R., Zhang, J., Steele, M.R., Romero, C.O., Kautzman, A.G., Schafer, D.P., and Vetter, M.L. (2019). Complement Targets Newborn Retinal Ganglion Cells for Phagocytic Elimination by Microglia. The Journal of neuroscience : the official journal of the Society for Neuroscience 39, 2025–2040. 10.1523/JNEUROSCI.1854-18.2018.

66. Woo, J.J., Pouget, J.G., Zai, C.C., and Kennedy, J.L. (2020). The complement system in schizophrenia: where are we now and what’s next? Molecular psychiatry 25, 114–130. 10.1038/s41380-019-0479-0.

67. Panagiotakopoulou, V., Ivanyuk, D., Cicco, S. de, Haq, W., Arsić, A., Yu, C., Messelodi, D., Oldrati, M., Schöndorf, D.C., and Perez, M.-J., et al. (2020). Interferon-γ signaling synergizes with LRRK2 in neurons and microglia derived from human induced pluripotent stem cells. Nature communications 11, 5163. 10.1038/s41467-020-18755-4.

68. Lee, Y., Ishikawa, T., Lee, H., Lee, B., Ryu, C., Davila Mejia, I., Kim, M., Lu, G., Hong, Y., and Feng, M., et al. (2025). Brain-wide mapping of immune receptors uncovers a neuromodulatory role of IL-17E and the receptor IL-17RB. Cell 188, 2203–2217.e17. 10.1016/j.cell.2025.03.006.

69. Mallick, R., Basak, S., Chowdhury, P., Bhowmik, P., Das, R.K., Banerjee, A., Paul, S., Pathak, S., and Duttaroy, A.K. (2025). Targeting Cytokine-Mediated Inflammation in Brain Disorders: Developing New Treatment Strategies. Pharmaceuticals (Basel, Switzerland) 18. 10.3390/ph18010104.

70. Martins, D., and Harrison, N.A. (2025). Cytokines as neuromodulators: insights from experimental studies in humans and non-human primates. Biological psychiatry. 10.1016/j.biopsych.2025.06.037.

71. Joy, M.T., Ben Assayag, E., Shabashov-Stone, D., Liraz-Zaltsman, S., Mazzitelli, J., Arenas, M., Abduljawad, N., Kliper, E., Korczyn, A.D., and Thareja, N.S., et al. (2019). CCR5 Is a Therapeutic Target for Recovery after Stroke and Traumatic Brain Injury. Cell 176, 1143–1157.e13. 10.1016/j.cell.2019.01.044.

72. Shen, Y., Zhou, M., Cai, D., Filho, D.A., Fernandes, G., Cai, Y., Sousa, A.F. de, Tian, M., Kim, N., and Lee, J., et al. (2022). CCR5 closes the temporal window for memory linking. Nature 606, 146–152. 10.1038/s41586-022-04783-1.

73. Cheadle, L., Rivera, S.A., Phelps, J.S., Ennis, K.A., Stevens, B., Burkly, L.C., Lee, W.-C.A., and Greenberg, M.E. (2020). Sensory Experience Engages Microglia to Shape Neural Connectivity through a Non-Phagocytic Mechanism. Neuron 108, 451–468.e9. 10.1016/j.neuron.2020.08.002.

74. Chen, Z.-P., Zhao, X., Wang, S., Cai, R., Liu, Q., Ye, H., Wang, M.-J., Peng, S.-Y., Xue, W.-X., and Zhang, Y.-X., et al. (2025). GABA-dependent microglial elimination of inhibitory synapses underlies neuronal hyperexcitability in epilepsy. Nature neuroscience 28, 1404–1417. 10.1038/s41593-025-01979-2.

75. Favuzzi, E., Huang, S., Saldi, G.A., Binan, L., Ibrahim, L.A., Fernández-Otero, M., Cao, Y., Zeine, A., Sefah, A., and Zheng, K., et al. (2021). GABA-receptive microglia selectively sculpt developing inhibitory circuits. Cell 184, 5686. 10.1016/j.cell.2021.10.009.

76. Wang, S., Hesen, R., Mossink, B., Nadif Kasri, N., and Schubert, D. (2023). Generation of glutamatergic/GABAergic neuronal co-cultures derived from human induced pluripotent stem cells for characterizing E/I balance in vitro. STAR protocols 4, 101967. 10.1016/j.xpro.2022.101967.

77. Savage, J.T., Ramirez, J.J., Risher, W.C., Wang, Y., Irala, D., and Eroglu, C. (2024). SynBot is an open-source image analysis software for automated quantification of synapses. Cell reports methods 4, 100861. 10.1016/j.crmeth.2024.100861.

78. Ewels, P.A., Peltzer, A., Fillinger, S., Patel, H., Alneberg, J., Wilm, A., Garcia, M.U., Di Tommaso, P., and Nahnsen, S. (2020). The nf-core framework for community-curated bioinformatics pipelines. Nature biotechnology 38, 276–278. 10.1038/s41587-020-0439-x.

79. Korsunsky, I., Millard, N., Fan, J., Slowikowski, K., Zhang, F., Wei, K., Baglaenko, Y., Brenner, M., Loh, P.-R., and Raychaudhuri, S. (2019). Fast, sensitive and accurate integration of single-cell data with Harmony. Nature methods 16, 1289–1296. 10.1038/s41592-019-0619-0.

