## Supplemental Figures for "MEA-LINK identifies the CCL4-CCR5 axis in neuronal hyperactivity control by human microglia"

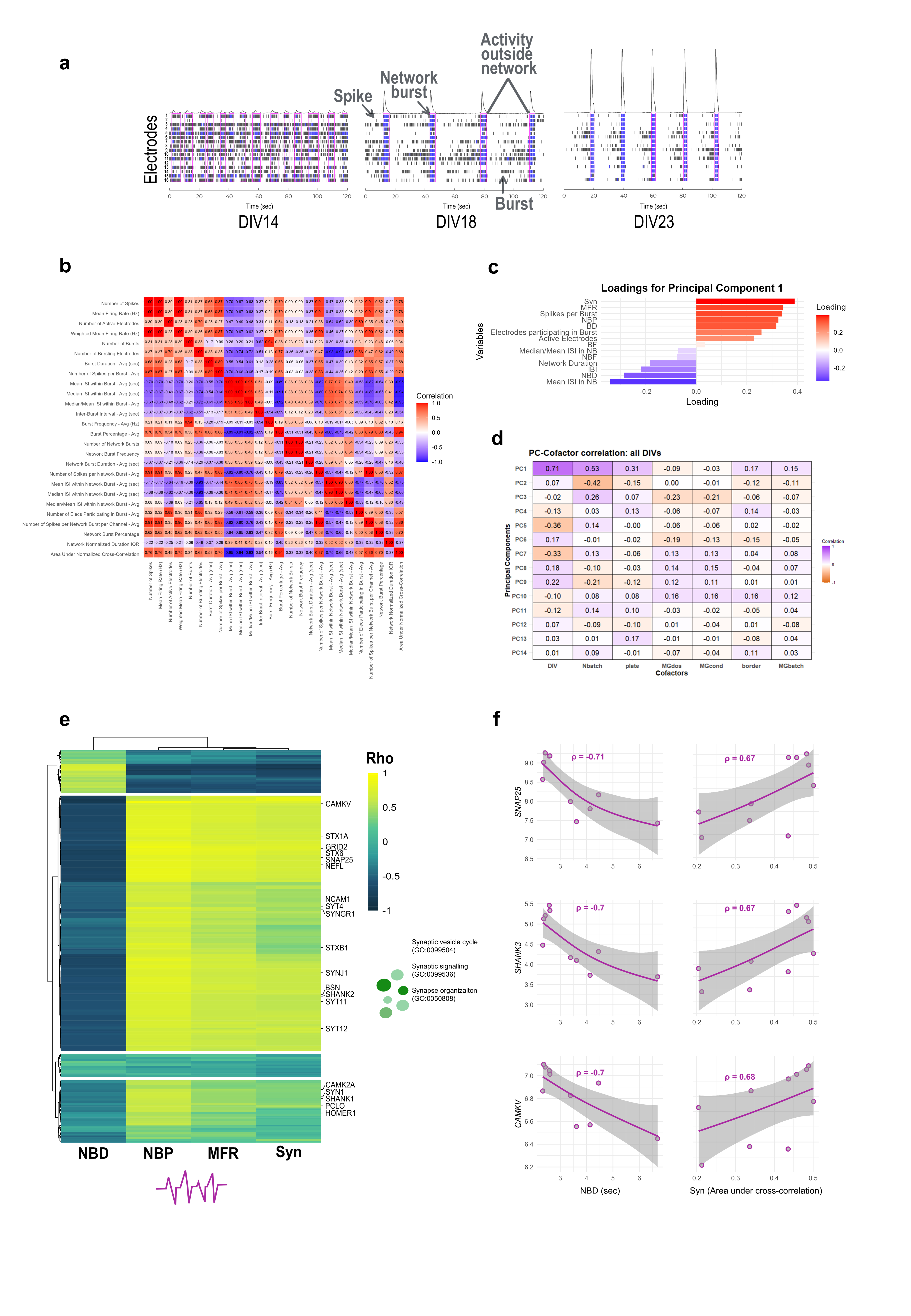

**Supplementary Figure 1.**

**(a)** Representative MEA activity raster plots during development showing increase in network organization with time. Spikes and Network bursts are annotated as an example of MEA analysis. **(b)** Correlation heatmap of all MEA parameters, N = 4, n = 1,071 neuronal network recordings. **(c)** Contributing MEA parameters to Principal Component (PC) 1 of all MEA recordings across development. **(d)** Correlation heatmap of PC 1 to 14 with co-variates in MEA experiments (DIV, batch, plate, MG-dosage, location of well on the plate). **(e)** MEA-RNA sequencing correlation heatmap using paired neuronal network activity measurements and transcriptomic profiles of the neuronal networks, N = 3, n = 10 neuronal networks. **(f)** Correlation plots of network burst duration and synchrony with *SNAP25*, *SHANK3*, and *CAMKV*, N = 3, n = 10 neuronal networks.

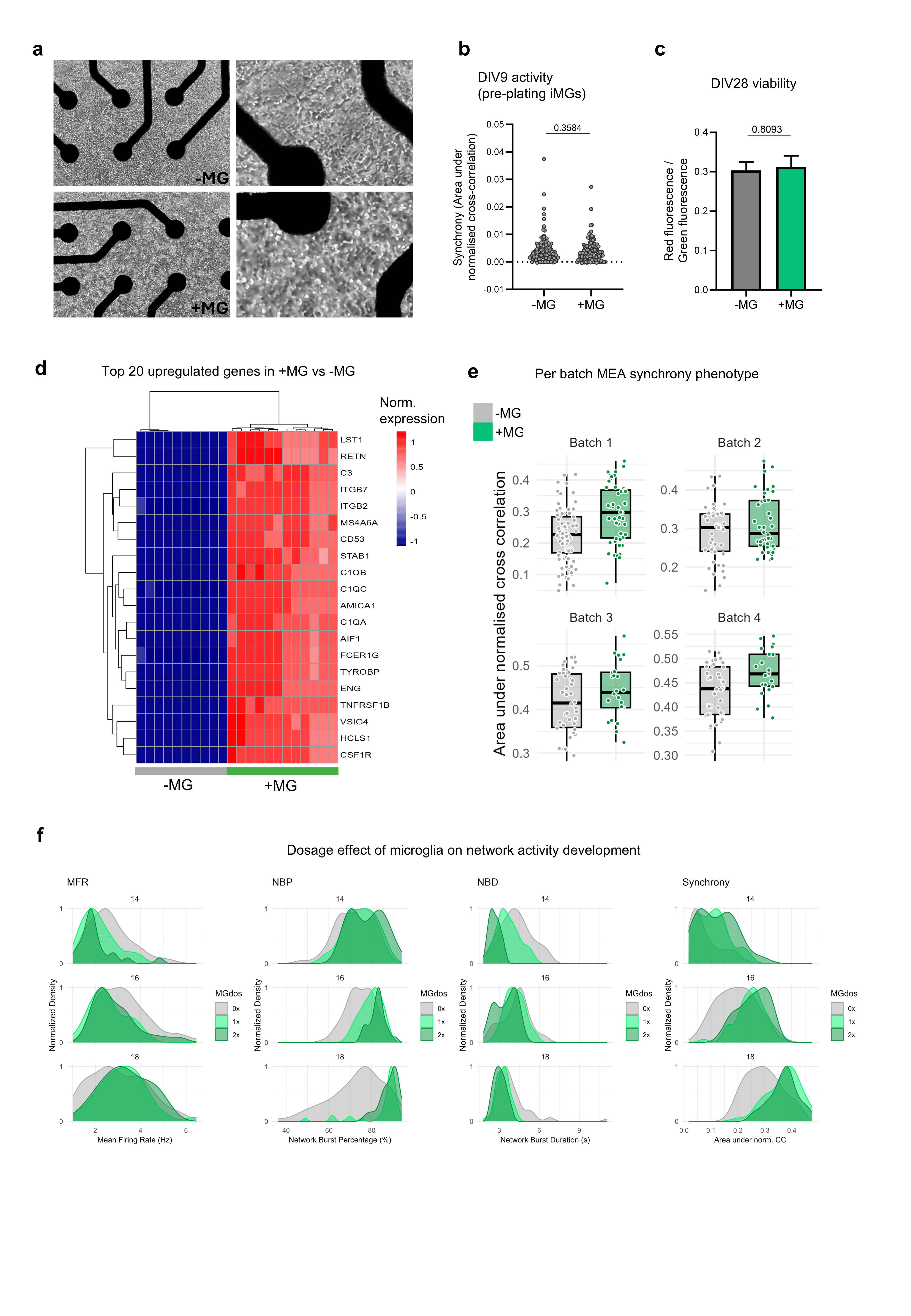

**Supplementary Figure 2.**

**(a)** Representative brightfield images of neuronal networks with and without microglia plated in MEA wells. **(b)** Synchrony at DIV9 before microglia were added to the neuronal networks, N = 4, n = 120 per condition, unpaired t-test. **(c)** Quantification of Live-Dead staining in neuronal networks at DIV28, N = 2, n = 9 per condition, unpaired t-test. **(d)** Heatmap of the top 20 upregulated genes in +MG networks compared to -MG networks, N = 3, n = 10 for -MG, 12 for +MG. **(e)** Synchrony quantification in neuronal networks shown for each of the four batches. **(f)** Density distribution plots of MFR, NBP, NBD, and synchrony at DIV14, 16, and 18 comparing neuronal networks with different microglial dosages: without microglia (0x), with 1:5 microglia (1x) and 2:5 microglia (2x), N = 2, n = 390 neuronal network recordings.

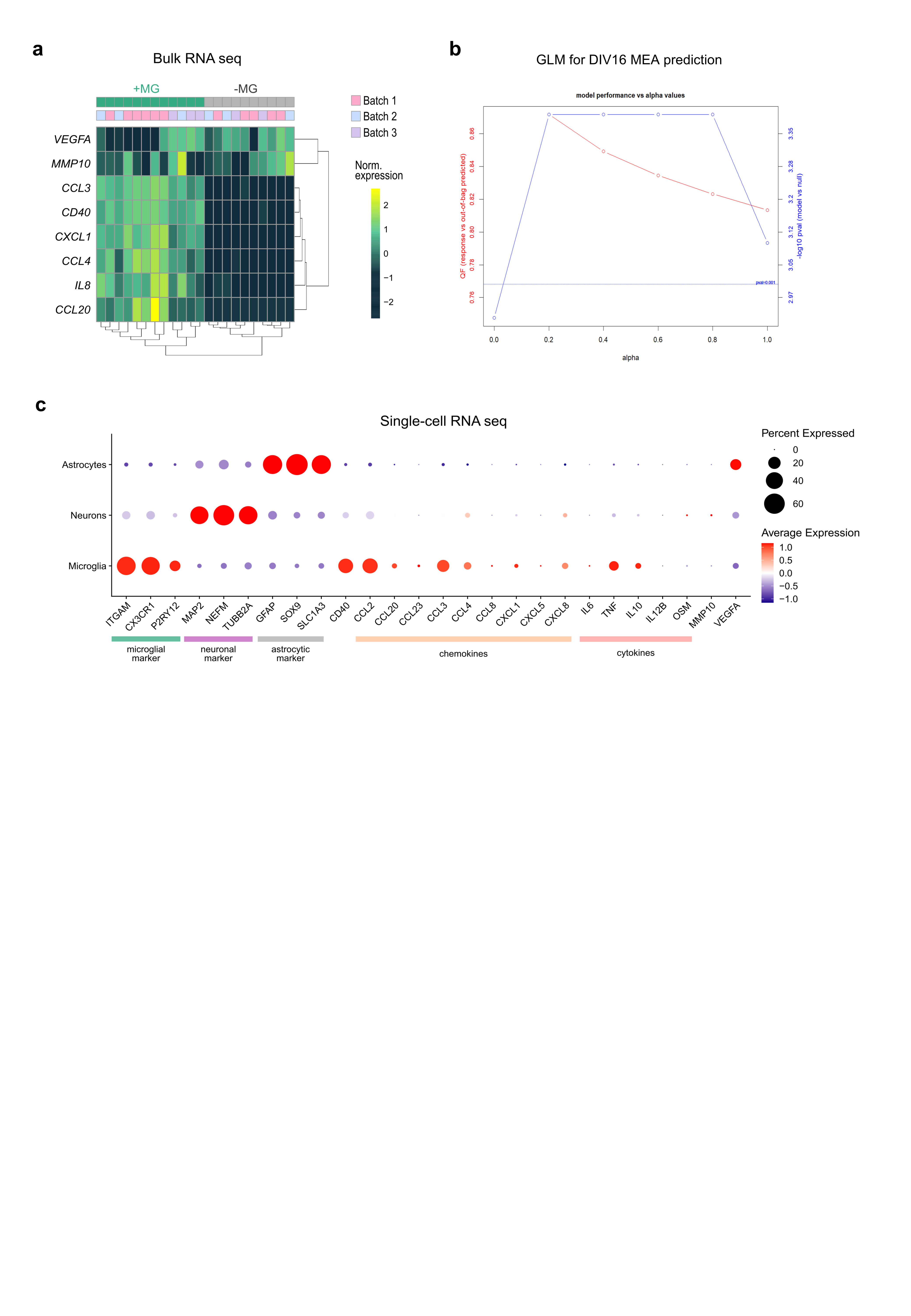

**Supplementary Figure 3.**

**(a)** Heatmap of RNA expression of candidates found with Olink proteomics in +MG and -MG neuronal networks, N = 3, n = 10 for -MG, 12 for +MG. **(b)** Performance comparison of generalised linear models using different alpha values, in -MG networks at DIV16, input = protein levels (NPX), output = network burst duration (sec). **(c)** Single-cell RNA sequencing data of cultures showing average gene expression of cell-type specific genes and candidates identified by Olink proteomics, N = 3, n = 22,355 single cells.

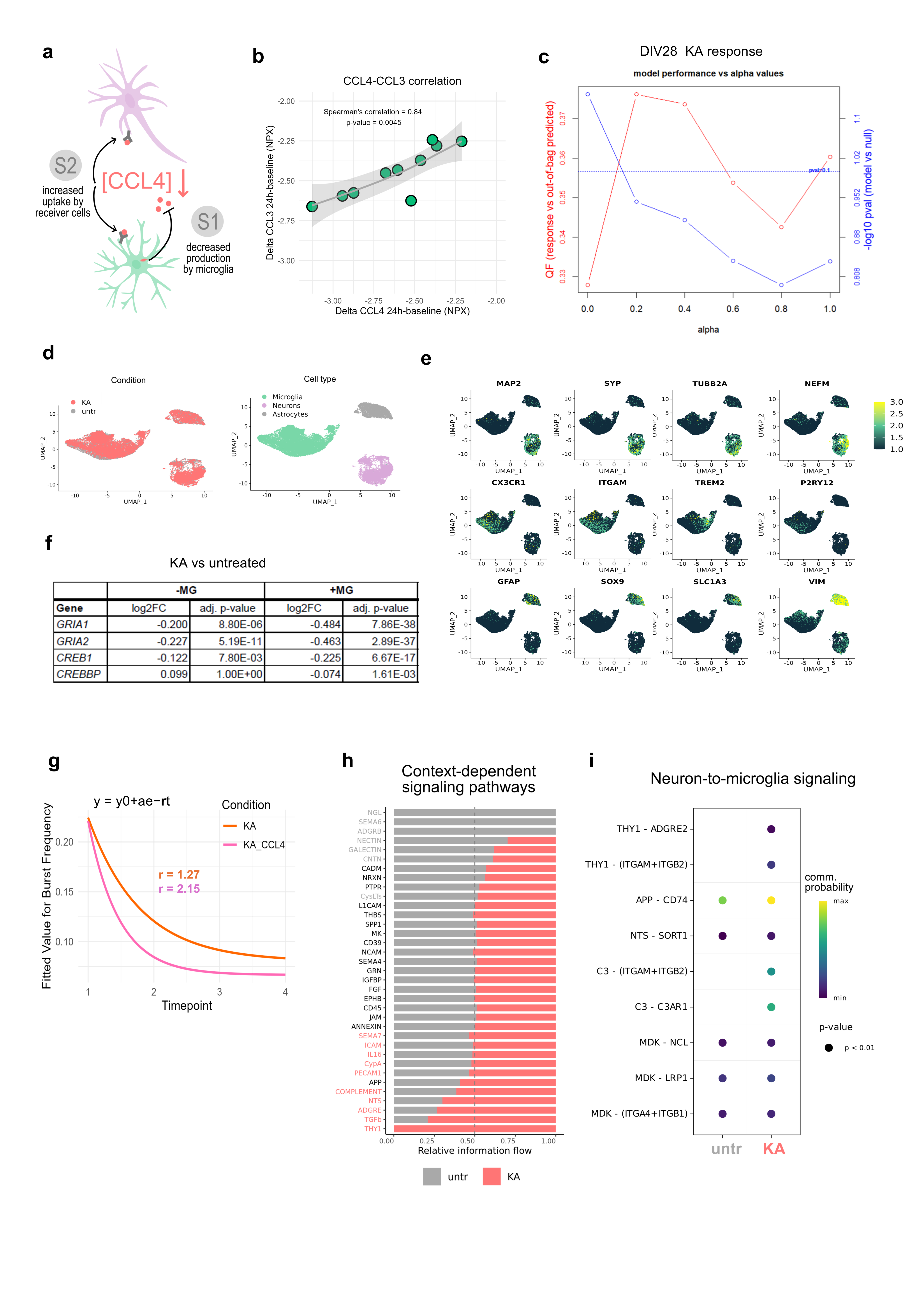

**Supplementary Figure 4.**

**(a)** Schematic overview of two possible scenarios for decrease in CCL4 concentration in the supernatant. Either by decreased production of microglia (S1) or increased uptake by receiver cells via CCR5 (S2). **(b)** Correlation plot of change in CCL4 and change in CCL3 (∆NPX) after KA treatment in +MG networks, n = 10. **(c)** Performance comparison of generalised linear models using different alpha values, in +MG networks at DIV28, input = change in protein levels after KA treatment (∆NPX), output = change in mean firing rate after KA treatment (∆MFR). **(d)** Uniform manifold approximation and projection (UMAP) analysis of single-cell RNA-seq data of cultures at DIV 28 showing distribution of KA treatment (left) and clustering of the three cell types (right), N = 3, n = 49,678 single cells (45% untreated, 55% KA). **(e)** UMAP analysis showing expression of cell-type specific markers for neurons (top), microglia (middle) and astrocytes (bottom). **(f)** Change in expression of *GRIA1*, *GRIA2*, *CREB1*, and *CREBBP2* in -MG and +MG networks after KA treatment. **(g)** Fitted exponential decay function on burst frequency recovery after KA-induced hyperactivity comparing addition of CCL4. **(h)** Predicted signaling pathways comparing untreated vs KA-treated cultures using CellChat. **(i)** Predicted ligand-receptor pairs involved in neuron-to-microglia signaling comparing untreated vs KA-treated cultures.
